# DNA topological regulation by topoisomerase IIβ-DNA-PK interaction is important for controlled hypoxia-inducible gene expression

**DOI:** 10.1101/2025.01.06.631366

**Authors:** Heeyoun Bunch, Jihye Park, Jasper K. Solverson, Jeongho Lee, Ji-Seung Yoo, Huiming Lu, Jaeyeon Jang, Young-Ran Yoon, Jae-Han Jeon, Matthew J. Schellenberg, Inuk Jung, Stuart K. Calderwood, Keunsoo Kang, Anthony J. Davis

**Affiliations:** School of Applied Biosciences, Kyungpook National University, Daegu 41566, Republic of Korea; Department of Applied Biosciences, Kyungpook National University, Daegu 41566, Republic of Korea; Department of Microbiology, College of Bio-convergence, Dankook University, Cheonan 31116, Republic of Korea; Department of Biochemistry and Molecular Biology, Mayo Clinic, Rochester, MN 55905, USA; School of Life Sciences, BK21 FOUR KNU Creative Bioresearch Group, Kyungpook National University, Daegu 41566, Republic of Korea; Department of Radiation Oncology, University of Texas Southwestern Medical Center, Dallas, TX 75390, USA; School of Computer Science and Engineering, Kyungpook National University, Daegu 41566, Republic of Korea; School of Medicine, Kyungpook National University, Daegu 41944, Republic of Korea; Department of Internal Medicine, Kyungpook National University Chilgok Hospital, School of Medicine, Kyungpook National University, Daegu 41404, Republic of Korea; Department of Radiation Oncology, Beth Israel Deaconess Medical Center, Harvard Medical School, Boston 02115, USA

**Keywords:** Hypoxia-inducible genes, topoisomerase IIβ, DNA-PK, RNA polymerase II transcription, gene regulation, DNA topology

## Abstract

Hypoxic stress responses are essential for cellular and organismal survival and drive gene regulation across diverse biological pathways, including cell cycle progression and energy metabolism. Here, we show that topoisomerase IIβ (TOP2B) regulates DNA topology and transcription of hypoxia-inducible genes (HIGs) in a DNA-dependent protein kinase (DNA-PK)-dependent manner. Integrated cellular, biochemical, and genomic analyses reveal an antagonistic yet correlated relationship between TOP2B and DNA-PK activities. Under normoxic conditions, TOP2B associates with HIGs and represses transcription by suppressing DNA negative supercoiling formation and accessibility. Under hypoxic exposure, TOP2B dissociates from HIGs, while DNA-PK and HIF1α are recruited to activate HIGs. Notably, DNA-PK is required for the repressive function of TOP2B, as DNA-PK knockout abrogates TOP2B release and activity, resulting in elevated expression of a number of HIGs. Mechanistically, DNA-PK phosphorylates TOP2B at T1403, stimulating its catalytic activity to restrain DNA unwinding and accessibility, thereby suppressing HIG transcription. Collectively, these findings identify a novel role for TOP2B and DNA-PK-mediated regulation of TOP2B as key transcriptional regulators of HIG expression. We propose that phosphorylation-dependent modulation of TOP2B activity is coordinated with transcriptional states and determines DNA topology to regulate Pol II transcription.

## INTRODUCTION

Transcription is the first step of gene expression and involves the synthesis of nascent RNA molecules catalyzed by RNA polymerases. The engagement, catalytic activity, and dissociation of RNA polymerases are tightly regulated by transcription factors and chromosome-associated elements^1^. In eukaryotes, RNA polymerase II (Pol II) is responsible for the transcription of messenger RNAs (mRNAs) and most of the small- and long-noncoding RNAs, excluding rRNAs and tRNAs. Since the discovery of Pol II in 1969, extensive research has elucidated the molecular mechanisms governing Pol II-mediated gene regulation^2^. Such mechanisms, in a majority of genes, occur throughout the stages of transcription of initiation, early elongation, processive elongation, and termination, in response to the signals that activate or repress gene expression^3,4^.

Gene regulation enables cells to respond to intrinsic metabolic demands, extracellular signals, and the maintenance of homeostasis. Hypoxia is an acute cellular stress that arises when oxygen availability is insufficient to meet metabolic requirements^5^. In mitochondria, oxygen serves as the terminal electron acceptor in the electron transport chain, where its reduction to H_2_O drives the generation of an electrochemical gradient across the mitochondrial inner membrane^6^. The gradient powers ATP synthase-mediated ATP production, the principal organismal chemical energy source^6^. Consequently, the optimal concentration of O_2_ is essential for efficient energy metabolism and sustaining cellular viability, function, and growth^7,8^. Under adverse conditions, such as infection, inflammation, cardiovascular diseases, and cancer, reduced oxygen availability imposes hypoxic stress, resulting in adaptive responses that include angiogenesis, metabolic shift to glycolysis, and enhanced migratory and metastatic capacity^9–12^. Activation of these pathways is mediated by a set of hypoxia-inducible genes (HIGs). The best characterized mechanism governing HIG activation involves hypoxia-inducible factor 1α (HIF1α), a transcription factor that is stabilized under low O_2_ conditions in cells^5,13–16^. Stabilized HIF1α translocates into the nucleus, heterodimerized with HIF1β, and binds hypoxia-responsive elements within target gene promoters to trigger transcriptional activation^8^.

The DNA-dependent protein kinase (DNA-PK) holoenzyme is a serine/threonine kinase composed of the DNA-PK catalytic subunit (hereafter DNA-PK) and the Ku70/Ku80 heterodimer. DNA-PK is well-known for its role in the repair of DNA double-stranded breaks^17,18^. The protein functions as a central factor for DNA repair via nonhomologous end joining, although some studies suggest that it also facilitates DNA repair via homologous recombination^18–20^. In our previous work, we unexpectedly identified that DNA-PK and one of its substrates, TRIM28^21^, in regulating the Pol II promoter-proximal pause (hereafter Pol II pausing) release at the *HSPA1B* (*HSP70*) and immediate early genes (IEGs), including *EGR1*, *JUN*, *FOS*, and *MYC*^22–25^. TRIM28 stabilizes Pol II pausing during the resting state of transcription, whereas upon transcriptional activation, it is phosphorylated at S824 by DNA-PK and ATM, an event that is required for effective Pol II pause release^22^. Notably, recent studies have extended these findings demonstrating that DNA-PK and TRIM28 also act as transcriptional activators of HIGs, regulating Pol II pausing and pause release at representative HIGs^26,27^ in a manner analogous to their roles in *HSP70* and IEG activation^23,24^. In both studies, inhibition of DNA-PK catalytic activity blocks substrate phosphorylation, which impairs gene activation. Collectively, these studies underscore a crucial role of DNA-PK as a regulator of transcriptional elongation at these genes.

Mammalian topoisomerase II (TOP2) is a key regulator of DNA topology, whose torsion-relieving catalysis is required for effective DNA metabolism processes, including replication, transcription, and DNA compaction into nucleosomes^28,29^. Recent studies from our group and others have demonstrated that topoisomerase IIβ (TOP2B) is recruited to the transcription start sites (TSSs) and plays a functional role in the transcriptional activation of estrogen- and androgen receptor genes, *HSP70*, neurotransmitter-inducible genes, retinoic acid responsive genes, and IEGs^23,30–37^. Although it remains unclear whether the DNA strand break (DSB) generated by TOP2B is programmed or accidental, the DNA break and damage response signaling caused by TOP2B catalytic activity stimulates the activation of these genes^23,36^. Importantly, the topological states of these genes before and during transcriptional activation remain to be elucidated. Excessive positive or negative supercoiling can impede Pol II progression; however, accumulating evidence suggests that negative supercoiling at promoters, generated by Pol II forward translocation, favors transcription, whereas positive supercoiling ahead of elongating Pol II is inhibitory to enzyme progression^38–43^. These findings raise the possibility that TOP2B exerts DNA topology-dependent biphasic effects; neutralization of promoter-associated negative supercoiling may suppress transcriptional activation, whereas relief of positive supercoiling in the downstream of Pol II may promote productive elongation. Regulation of TOP2B binding, catalytic activity, and dissociation during transcriptional activation is thought to be regulated via post-translational modifications. For instance, our previous studies showed that TOP2B ubiquitination by the BRCA1–BARD1 complex enhances its DNA-binding affinity, while TOP2B phosphorylation by ERK2 modulates TOP2B catalytic activity toward positive and negative DNA supercoils and TOP2B dissociation from its target genes^35,36^.

Although both TOP2B and DNA-PK have been implicated in the activation of estrogen- and androgen receptor genes, *HSP70*, neurotransmitter- and retinoic acid-inducible genes, and IEGs, and DNA-PK has been reported to play an important role in hypoxic gene regulation^23,27,30,31,33–37^, the contribution of TOP2B in HIG transcription has not been previously examined. Notably, TOP2B physically interacts with Ku70, whereas DNA-PK neither binds to nor is activated by the broken DNA end of the TOP2B-DNA adduct^44,45^. These observations raise the intriguing possibility of functional crosstalk and regulation between TOP2B catalytic activity and DNA-PK signaling during HIG transcription. In this study, we investigated whether TOP2B participates in HIG transcriptional regulation, whether it functionally interacts with DNA-PK, and how this interaction influences transcriptional output. We found that TOP2B represses HIG transcription under normoxic conditions and becomes inactivated and dissociates from the TSSs of HIG genes upon hypoxic stress. DNA-PK exerts biphasic effects on HIG expression, as DNA-PK knockout (KO) or catalytic inactivation differentially increases or decreases distinct subsets of HIGs. Mechanistically, DNA-PK promotes the repressive activity of TOP2B and regulates its catalytic activity and release from chromatin. In the absence of DNA-PK, TOP2B occupancy at HIG loci increases and transcription is stimulated. Conversely, TOP2B limits the recruitment of DNA-PK and HIF1α at HIGs and attenuates DNA-PK phosphorylation at T2609, and suppresses HIG transcription. Loss of TOP2B results in increased negative DNA supercoiling and chromatin openness at HIGs and across the genome. Notably, TOP2B KO enhances chromatin accessibility at HIGs, while reducing accessibility genome-wide, indicating locus-specific and global roles for TOP2B in regulating DNA topology. Finally, mutational and topological analyses reveal that DNA-PK phosphorylates TOP2B at its carboxyl-terminal domain (CTD) including at T1403 and that this post-translational modification is important for TOP2B-mediated suppression of HIG transcription through the removal of negative supercoiling.

## RESULTS

### HIG expression is regulated by TOP2

DNA structure and topology have been recognized as critical transcription regulatory elements of diverse stimulus-inducible metazoan genes^30,33,34,36,46–52^. In humans, TOP1 and TOP2 resolve the negative and positive DNA supercoiling associated with DNA metabolism, including transcription^53^. In this study, we have investigated whether human TOP2 is involved in transcriptional regulation of HIGs. Representative HIGs, including *p21*, *ALDOC*, *GLUT*s, and *IL1β*, were examined throughout the study^5,11,14,54–56^. To induce hypoxic responses, human neuroblastoma cells were treated with CoCl_2_, a chemical validated to mimic hypoxic conditions in multiple studies^12,57,58^, at a final concentration of 200 µM for 12 h. Quantitative RT-PCR (qRT-PCR) analysis revealed significant transcriptional activation at *p21*, *ALDOC*, *IL1β*, and *GLUT1* following CoCl_2_ treatment (**Fig. 1A** and **Supplementary Figure 1A**). Under the same hypoxic conditions, chromatin immunoprecipitation (ChIP)-PCR analysis showed that the master transcriptional activator for HIGs, HIF1α was recruited to the TSSs of these genes, whereas the level of TOP2B occupancy was notably decreased in the TSSs as well as the gene bodies (GBs) (**Fig. 1B** and **Supplementary Figure 1B**). Although TOP2A has traditionally been regarded primarily as a DNA replication factor, recent studies have implicated TOP2A in transcription regulation^59,60^. Therefore, we assessed TOP2A occupancy at the representative HIGs and observed a pattern similar to that of TOP2B, with both enzymes dissociating from HIG promoters upon transcriptional activation (**Fig. 1C**). Notably, TOP2B is generally known as a transcriptional activator that is recruited to stimulated stress-inducible genes, including *HSP70*, IEGs, estrogen- and androgen activated genes, and neurotransmitter-inducible genes^23,24,30,33–36,61^. Thus, the coordinated release of both TOP2A and TOP2B from representative HIG loci during transcription activation represents an unexpected mode of regulation, suggesting a distinctive role for TOP2 enzymes in HIG transcription.

**Fig. 1.**
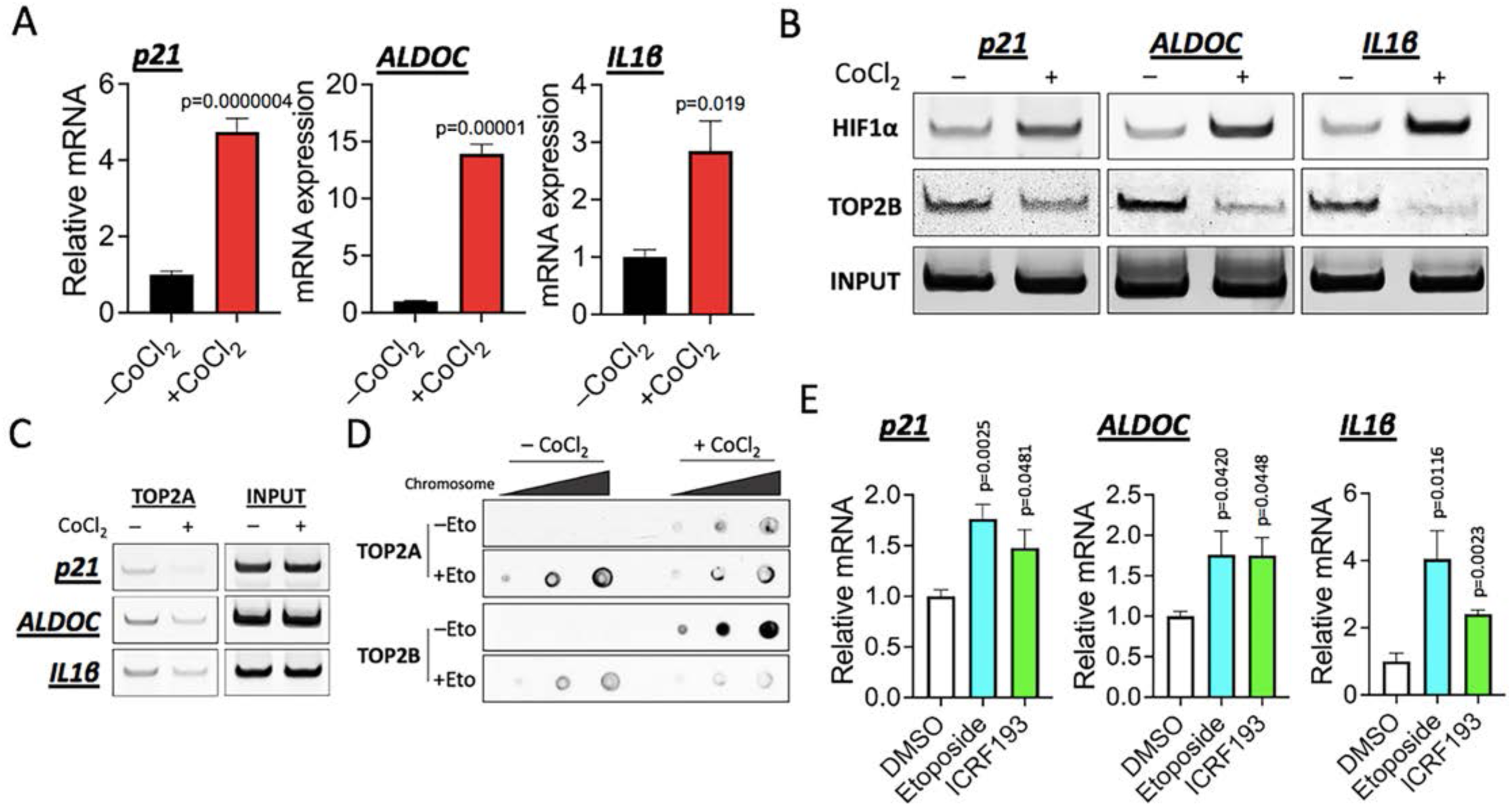
HIG expression is regulated by TOP2. (**A**) qRT-PCR results presenting the effect of hypoxic-stress caused by CoCl_2_ treatment (+CoCl_2_, throughout the figures) on the transcription of representative HIGs, *p21*, *ALDOC*, and *IL1β*, in SH-SY5Y cells (n= 3 biologically independent samples). An equal amount of H_2_O, the solvent for CoCl_2_, was applied as the control (–CoCl_2_, throughout the figures). Data are presented as mean values and SD. *P*-values for the bar graphs were calculated with the unpaired, one-sided Student’s t-test. (**B)** Representative ChIP-PCR results showing HIF1α and TOP2B occupancies on the representative HIGs, *p21*, *ALDOC*, and *IL1 β* under normoxia and hypoxia in SH-SY5Y cells. (**C**) Representative ChIP-PCR results showing TOP2A occupancy on the representative HIGs, *p21*, *ALDOC*, and *IL1 β* under normoxia and hypoxia in SH-SY5Y cells. (**D**) Representative ICE assay results showing the TOP2A and TOP2B proteins engaged with the chromosomal DNA under normoxia and hypoxia, with or without etoposide treatment (+Eto, 50 μM, 1 h; –Eto, DMSO). Three dots per –CoCl_2_ or +CoCl_2_ set are 22, 44, and 66 ng genomic DNA (chromosome) loaded onto nitrocellulose membrane, followed by immunoblotting using TOP2A or TOP2B antibody. (**E**) qRT-PCR results indicating that under hypoxic stresses, inhibiting TOP2B using etoposide and ICRF193 stimulates transcription at the representative HIGs, *p21*, *ALDOC*, and *IL1 β* (n= 3 biologically independent samples). Data are presented as mean values and SEM. *P*-values for the bar graphs were obtained through one-way ANOVA with multiple comparisons.

To understand TOP2B–genome interactions under hypoxic conditions, we next assessed TOP2 engagement with chromosomal DNA using an *in vivo* complex of enzyme (ICE) assay. CoCl_2_-induced hypoxia increased the levels of both TOP2A and TOP2B associated with the genome (**Fig. 1D**). As expected, treatment with etoposide, a TOP2 poison that traps the proteins in the TOP2–DNA covalent cleavage complex (TOP2cc)^62^, led to increased association of both TOP2A and TOP2B with DNA under normoxic conditions. Interestingly, however, under hypoxic conditions, the level of etoposide-trapped TOP2B protein was markedly reduced (**Fig. 1D** and **Supplementary Figure 1C**), indicating either a reduction in catalytic activity of TOP2B, enhanced removal of TOP2cc, or a combination of both mechanisms under hypoxia. In our previous studies, we showed that etoposide activates, whereas ICRF193, a catalytic inhibitor of TOP2, represses transcription of IEGs, such as *EGR1* and *FOS*^23,36^. To assess whether TOP2 activity similarly influences HIG expression, SH-SY5Y cells were treated with etoposide (10 µM) or ICRF193 (10 μM) in the presence of CoCl_2_ (200 μM) for 3 h, followed by qRT-PCR analysis. Interestingly, both inhibitors increased mRNA levels of representative HIGs (**Fig. 1E**), indicating that TOP2 catalytic activity restrains productive transcription. Collectively, the data presented in **Fig. 1** uncover a novel, repressive role for TOP2 in regulating the transcription of HIGs through some unique mechanisms.

### DNA-PK interacts with and regulates TOP2B in HIGs and genome-wide

As noted above, DNA-PK has been implicated in HIG transcription^27^, and the Ku70 subunit of DNA-PK holoenzyme has been reported to physically interacts with TOP2^44^. We therefore examined whether TOP2B also interacts with DNA-PK. To assess protein-protein interactions, immunoprecipitation assays were performed using HeLa nuclear extracts. To restrict detection to stable protein interactions, immunoprecipitates were subjected to highly stringent wash conditions ( 0.15–0.5 M HEGN with >130-fold bead volume), prior to immunoblotting analysis. Under these conditions, TOP2B robustly co-precipitated DNA-PK, and reciprocal immunoprecipitation confirmed this interaction (**Fig. 2A**), indicating a strong and stable physical interaction between TOP2B and DNA-PK. Consistent with a previously report^27^, HIF1α was also found to interact with DNA-PK in these assays. In contrast, no interaction between TOP2B and HIF1α was detected (**Fig. 2A**).

**Fig. 2.**
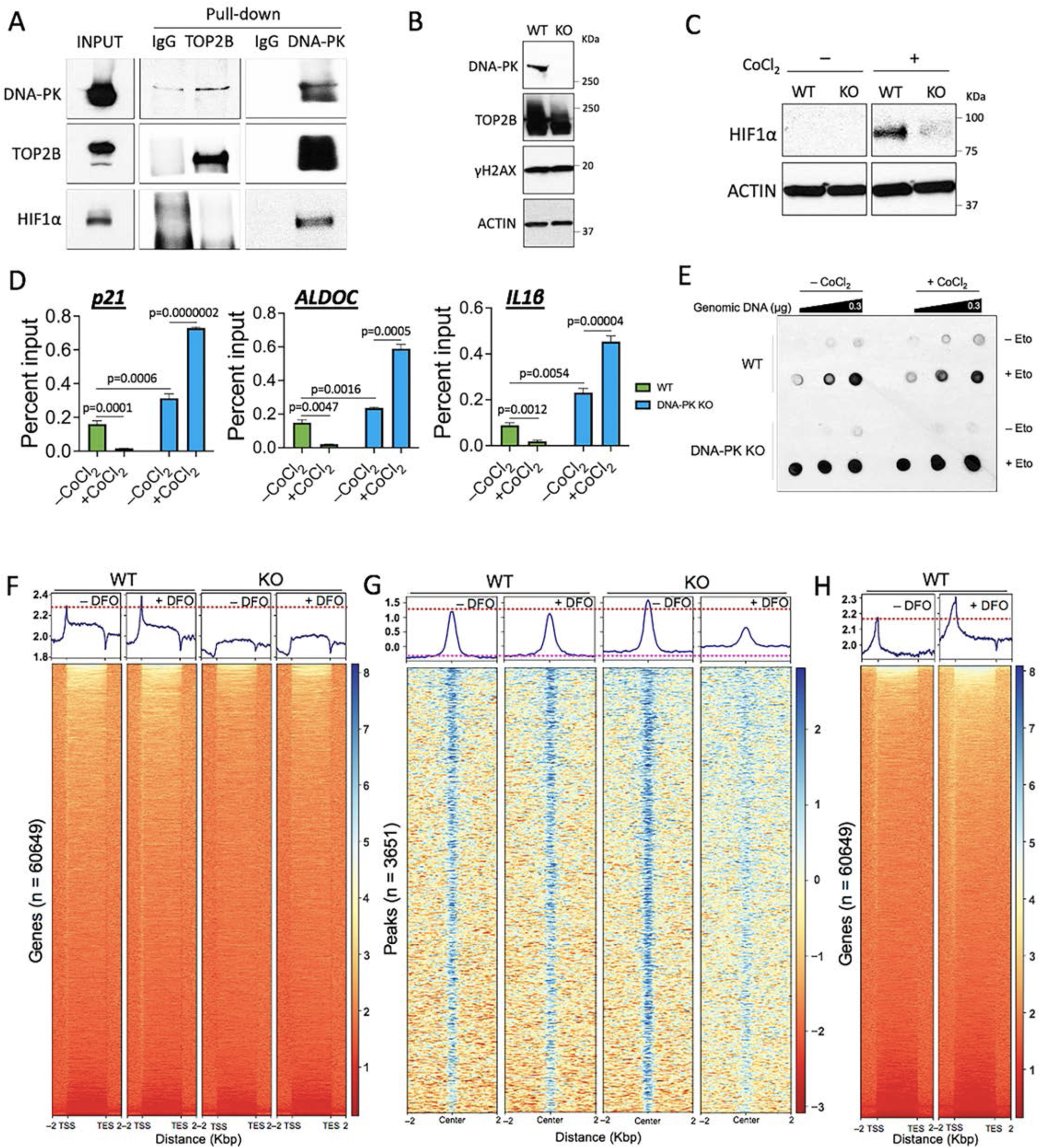
DNA-PK interacts with and regulates TOP2B at HIGs and genome-wide. (**A**) Immunoprecipitation results showing TOP2B–DNA-PK interaction. INPUT, 0.7% HeLa NE used for each reaction. (**B**) Immunoblots of DNA-PK, TOP2B, and γH2AX, comparing WT and DNA-PK KO HCT116 cells. ACTIN, a reference. (**C**) Immunoblots of HIF1α, comparing WT and DNA-PK KO cells. ACTIN, a reference. (**D**) ChIP-qPCR results showing TOP2B occupancy changes in representative HIGs with or without DNA-PK (n = 3 independent experiments). Data are presented as mean values and SD. *P*-values for the bar graphs were calculated with the unpaired, one-sided Student’s t-test. (**E**) Representative ICE assay results showing the TOP2B proteins engaged with the chromosomal DNA under normoxia and hypoxia for 12 h, with or without etoposide treatment (+Eto, 50 μM, 1 h; – Eto, DMSO). Three dots per –CoCl_2_ or +CoCl_2_ set are 75, 150, and 300 ng genomic DNA (chromosome) loaded onto nitrocellulose membrane, followed by immunoblotting using TOP2B antibody. (**F**) ChIP-seq data showing HIF1α occupancies in all genes (n = 60649) under normoxia (– DFO) or hypoxia (+ DFO) in WT and DNA-PK KO HCT116 cells (n = 2 independent experiments). (**G**) ChIP-seq data showing TOP2B peak changes (n = 3651) under normoxia (– DFO) or hypoxia (+ DFO) in WT and DNA-PK KO (KO) HCT116 cells (n = 2 independent experiments). (**H**) ChIP-seq data showing DNA-PK occupancies in in all genes (n = 60649) under normoxia (– DFO) or hypoxia (+ DFO) in WT cells (n = 2 independent experiments).

Next, we aimed to determine the functional significance of TOP2B-DNA-PK interaction in HIG transcription. To this end, we employed DNA-PK-KO HCT116 cells, which were previously generated and well-characterized, and we validated loss of DNA-PK expression by immunoblotting (**Fig. 2B**)^21,63^. Despite DNA-PK deletion, total TOP2B protein levels were largely unchanged, although a reduction in a slower-migrating, smeared TOP2B band, likely corresponding to post-translationally modified form, was observed (**Fig. 2B**). In addition, basal levels of γH2AX were not significantly affected by DNA-PK loss (**Fig. 2B**), consistent with previous reports showing that hypoxic stresses did not globally elevate γH2AX levels^27^ (**Supplementary Figure 2A**). Strikingly, loss of DNA-PK dramatically impaired stabilization of HIF1α under CoCl_2_-induced hypoxic conditions (**Fig. 2C** and **Supplementary Figure 2A**), indicating that DNA-PK modulates the HIF1α-mediated hypoxic responses.

To assess whether DNA-PK influences TOP2B chromatin association at HIGs, TOP2B occupancy at representative HIGs was examined by ChIP-qPCR in DNA-PK wild-type (WT) and KO cells under normoxic and hypoxic conditions. Strikingly, loss of DNA-PK significantly increased TOP2B occupancy at all three HIGs examined, *p21*, *ALDOC*, and *IL1β*, under normoxia (**Fig. 2D**). Moreover, the hypoxia-induced dissociation of TOP2B observed in WT cells was abolished in DNA-PK KO cells; instead, TOP2B occupancy further increased under hypoxic conditions in the absence of DNA-PK (**Fig. 2D**). Overall TOP2B catalytic activity on the chromosomal DNA also markedly increased in DNA-PK KO cells, under normoxia and hypoxia (**Fig. 2E** and **Supplementary Figure 2B**). To validate and extend these findings, we performed ChIP-seq analyses. WT and DNA-PK KO cells were treated with deferoxamine (DFO), an iron-chelating hypoxia-mimetic agent^64^, at a final concentration of 60 μM for 24 h. Consistent with hypoxia-induced transcriptional activation (**Fig. 2C**), HIF1α occupancy in the TSSs increased following DFO treatment in WT cells (**Fig. 2F**). As anticipated based on impaired HIF1α stabilization (**Fig. 2C**), genome-wide HIF1α binding was markedly reduced in DNA-PK KO cells under both normoxic and hypoxic conditions (**Fig. 2F** and **Supplementary Figure 2C**). TOP2B ChIP-seq analyses indicated a modest overall reduction in TOP2B genomic occupancy in WT cells under hypoxic stress (**Fig. 2G**). In contrast, deletion of DNA-PK increased TOP2B peak intensities under normoxic conditions (**Fig. 2G**), consistent with the ChIP-qPCR results (**Fig. 2D**). Strikingly, in the absence of DNA-PK, TOP2B occupancy became broadly elevated and less focal across the genome, with a particular pronounced increase under hypoxic conditions (**Fig. 2G**). These genome-wide changes indicate a critical role for DNA-PK in regulating TOP2B chromatin association and genomic distribution. Additionally, ChIP-seq analysis of DNA-PK itself showed increased occupancy at genomic loci under hypoxic stress, mirroring the recruitment pattern observed for HIF1α (**Fig. 2H** and **Supplementary Figure 2D**).

### HIG expression is regulated by the TOP2B–DNA-PK interaction

To determine how DNA-PK influences HIG expression, mRNA expression of representative HIGs in WT and DNA-PK KO HCT116 cells was quantified using qRT-PCR. Strikingly, DNA-PK KO resulted in significantly increased mRNA production at representative HIGs (**Fig. 3A**). This phenotype is due to DNA-PK enzymatic activity as similar results were observed in HCT116 cells with a mutation (K3753R) that inactivates the kinase activity of DNA-PK (kinase-dead, KD)**(Supplementary Figure 3A**)^63^. These results, together with the increased TOP2B occupancy observed in the absence of DNA-PK, suggest that DNA-PK promotes a repressive function of TOP2B at HIGs and that loss of DNA-PK permits TOP2B to support transcriptional activation (**Figs. 2D–H,3A** and **Supplementary Figure 3A**). Unexpectedly, enhanced HIG expression occurred despite impaired HIF1α stabilization in DNA-PK KO cells. To resolve this apparent discrepancy, we performed RNA-seq in DNA-PK WT and KO cells under normoxic conditions and following CoCl_2_-induced hypoxic stress (200 µM CoCl_2_ supplemented for 12 h). Comparative analysis identified a total of 258 genes with significantly altered expression under hypoxia [|Log_2_Fold change (FC)| > 0.5, p < 0.05], of which 192 were upregulated and 66 were downregulated (**Fig. 3B** and **Supplementary Data 1**). More broadly, loss of DNA-PK affected the expression of 4700 genes genome-wide (**Fig. 3C** and **Supplementary Data 2**), with approximately 58% (n = 2706 genes) showing increased expression in DNA-PK KO cells (**Fig. 3C** and **Supplementary Data 2)**. Focusing specifically on hypoxia-induced genes, 40.6% (n = 78 genes) were differentially expressed in DNA-PK KO cells (p-value of approximately 59.9% of HIGs < 0.05, thus considerable), with the majority (69 genes; 88.5%) exhibiting elevated expression without DNA-PK (**Fig. 3D** and **Supplementary Data 1,2**). These findings support a predominantly repressive role for DNA-PK in regulating HIG transcription. Examining individual HIGs with well-established roles in hypoxic responses revealed distinctive expression patterns depending on DNA-PK status (|FC| > 0.5, p-value < 0.05; **Fig. 3E**). Whereas a minority of genes (11.5%) including *MMP25*^65^ and *HK2*^66^ were downregulated, a larger subsets, including *ANKRD37*^67^, *DDIT4*^68^, *p21*, *ALDOC*, and *GLUT3*, displayed increased gene expression in DNA-PK KO cells (**Fig. 3E**).

**Fig. 3.**
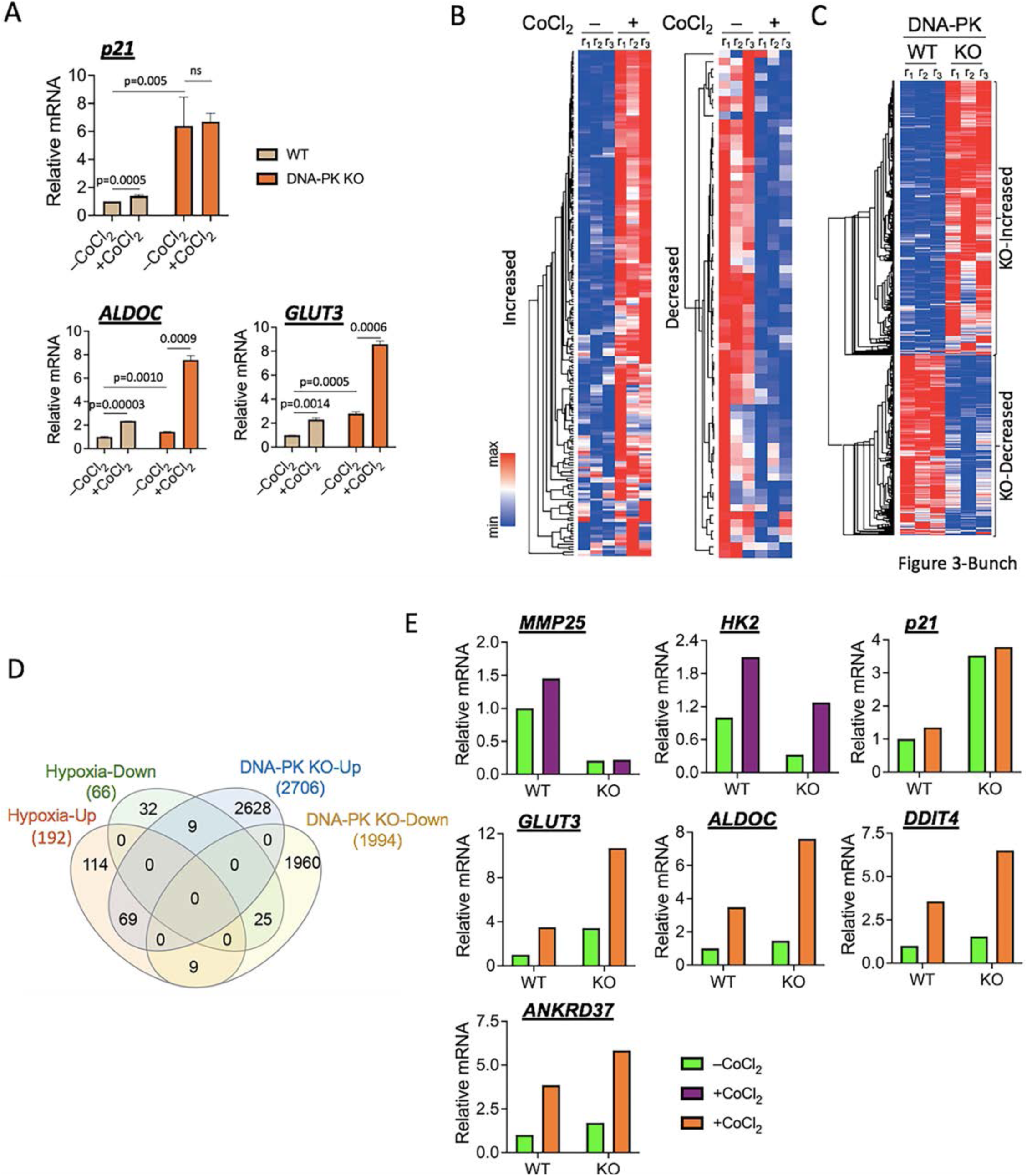
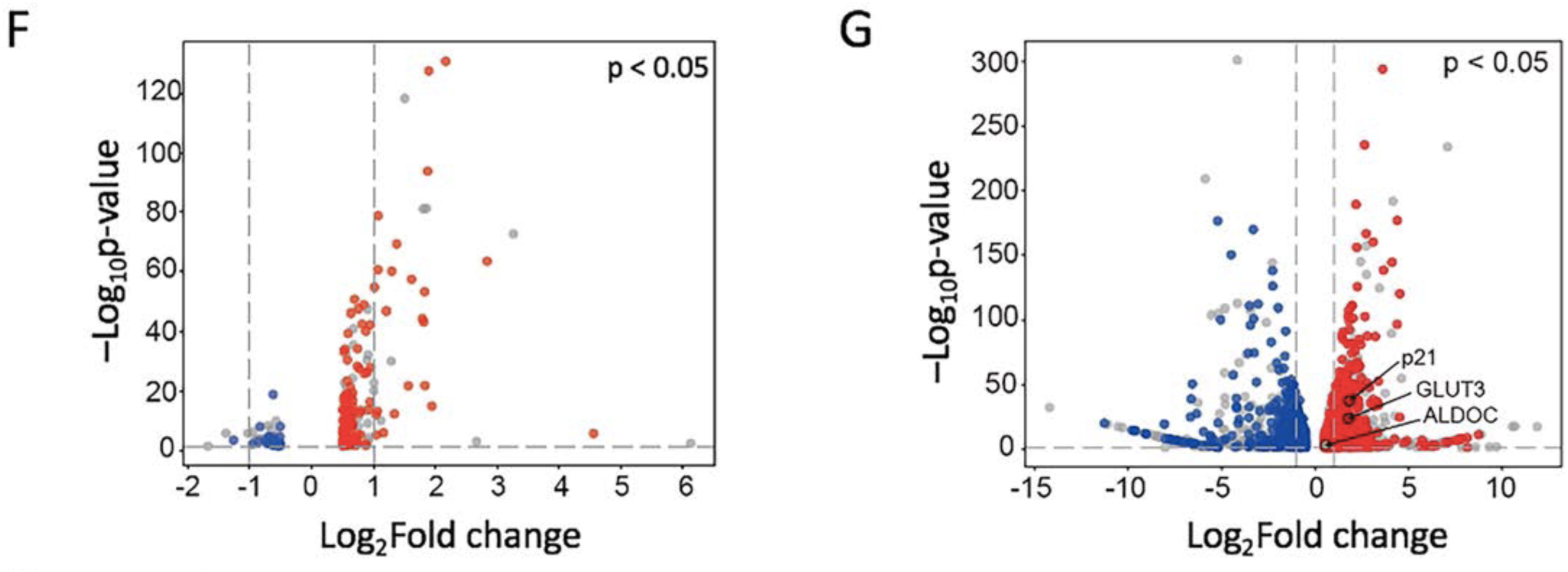
HIG expression is regulated by the TOP2B–DNA-PK interaction. (**A**) qRT-PCR results presenting the effects of DNA-PK KO on representative HIG transcription (n = 3). Data are presented as mean values and SD. *P*-values for the bar graphs were calculated with the unpaired, one-sided Student’s t-test. (**B**) Heatmaps illustrating HIG expression analyzed by RNA-seq data. Genes, whose expression is significantly increased (left, n = 192) or decreased (right, n = 66), are shown. |Log_2_FC| > 0.5, p < 0.05. Biological triplicates are shown as r_1–3_ (n = 3) ) in all RNA-seq data heatmaps of this study. (**C**) Heatmap presenting differentially expressed genes in the absence of DNA-PK. Upregulated (n = 2706) and downregulated (n = 1994) genes. |Log_2_FC| > 0.5, p < 0.05. (**D**) Venn diagram displaying the genes that are regulated by hypoxic stress and DNA-PK. (**E**) Graphs showing mRNA expression of important HIGs in RNA-seq data. Green-purple and green-orange graphs are the representative genes that are downregulated and upregulated in DNA-PK KO cells, respectively. p < 0.05. (**F**) Volcano plot exhibiting the genes that are occupied by TOP2B (red dots in upregulated genes or blue dots in downregulated genes under hypoxic stress). (**G**) Volcano plot exhibiting the genes that are occupied by TOP2B (red dots in upregulated genes or blue dots in downregulated genes in DNA-PK KO cells).

We also examined TOP2B occupancies at HIGs using the previous TOP2B ChIP-seq data (GSM2442946, **Supplementary Data 3**). Among the 192 identified HIGs, 76% (n = 146) were occupied by TOP2B (**Fig. 3F**, red dots). Approximately 42% of hypoxia-downregulated genes were bound by TOP2B (**Fig. 3F**, blue dots). Overall, more than 67% of hypoxia-regulated genes, whether upregulated or downregulated, exhibited TOP2B occupancy, indicating a broad involvement of TOP2B in transcriptional responses to hypoxia. We further analyzed TOP2B occupancy in DNA-PK-regulated genes. Approximately 54% of these genes upregulated (n = 1455, red dots), including *p21*, *GLUT3*, and *ALDOC*, and 48% downregulated genes (n = 957, blue dots) were bound by TOP2B (**Fig. 3G**). These findings suggested functional overlap between DNA-PK and TOP2B in the regulation of gene expression in humans. GO analyses of DNA-PK KO-induced, TOP2B-occupied genes revealed that enrichment in pathways related to control cellular morphogenesis and mobility (**Supplementary Figure 3B** and **Supplementary Data 4**), suggesting a suppressive role of DNA-PK and DNA-PK-TOP2B interactions on these genes. In contrast, GO analyses of DNA-PK KO-repressed, TOP2B-bound genes showed enrichment for ribonucleoprotein complex biogenesis, transcription, and DNA metabolism (**Supplementary Figure 3C** and **Supplementary Data 4**), suggesting the activating role of DNA-PK and DNA-PK-TOP2B interactions in these pathways. Collectively, these data presented in **Fig. 3** support a model, in which DNA-PK modulates TOP2B-dependent transcriptional regulation. Specifically, DNA-PK appears to facilitate TOP2B-mediated repression of HIG transcription, while promoting transcription of genes involved in RNA processing and genome maintenance, highlighting biphasic roles for TOP2B–DNA-PK interaction in global gene regulation.

### TOP2B regulates HIF1α and DNA-PK recruitment and activity at HIGs

Our data showed that under hypoxic stresses, TOP2B was released from representative HIGs in WT cells, whereas it was recruited to these genes in DNA-PK KO cells (**Figs. 1A,B,2D–H**), indicating that DNA-PK modulates TOP2B behavior and function. To further define this regulatory relationship, we examined DNA-PK regulation in TOP2B KO SH-SY5Y cells^37^. TOP2B KO was confirmed by immunoblotting (**Fig. 4A**). In TOP2B KO cells, total cellular DNA-PK protein levels were not noticeably changed (**Fig. 4A**), indicating that TOP2B does not regulate DNA-PK expression or protein stability. qRT-PCR analysis with the cells treated with 200 µM CoCl_2_ or H_2_O (–CoCl_2_, control) for 6 h showed that, with the except of *p21*, representative HIGs were transcriptionally activated under normoxic conditions in TOP2B KO cells and further induced under hypoxia (**Fig. 4B**). These findings confirm a suppressive role of TOP2B for HIG transcription.

**Fig. 4.**
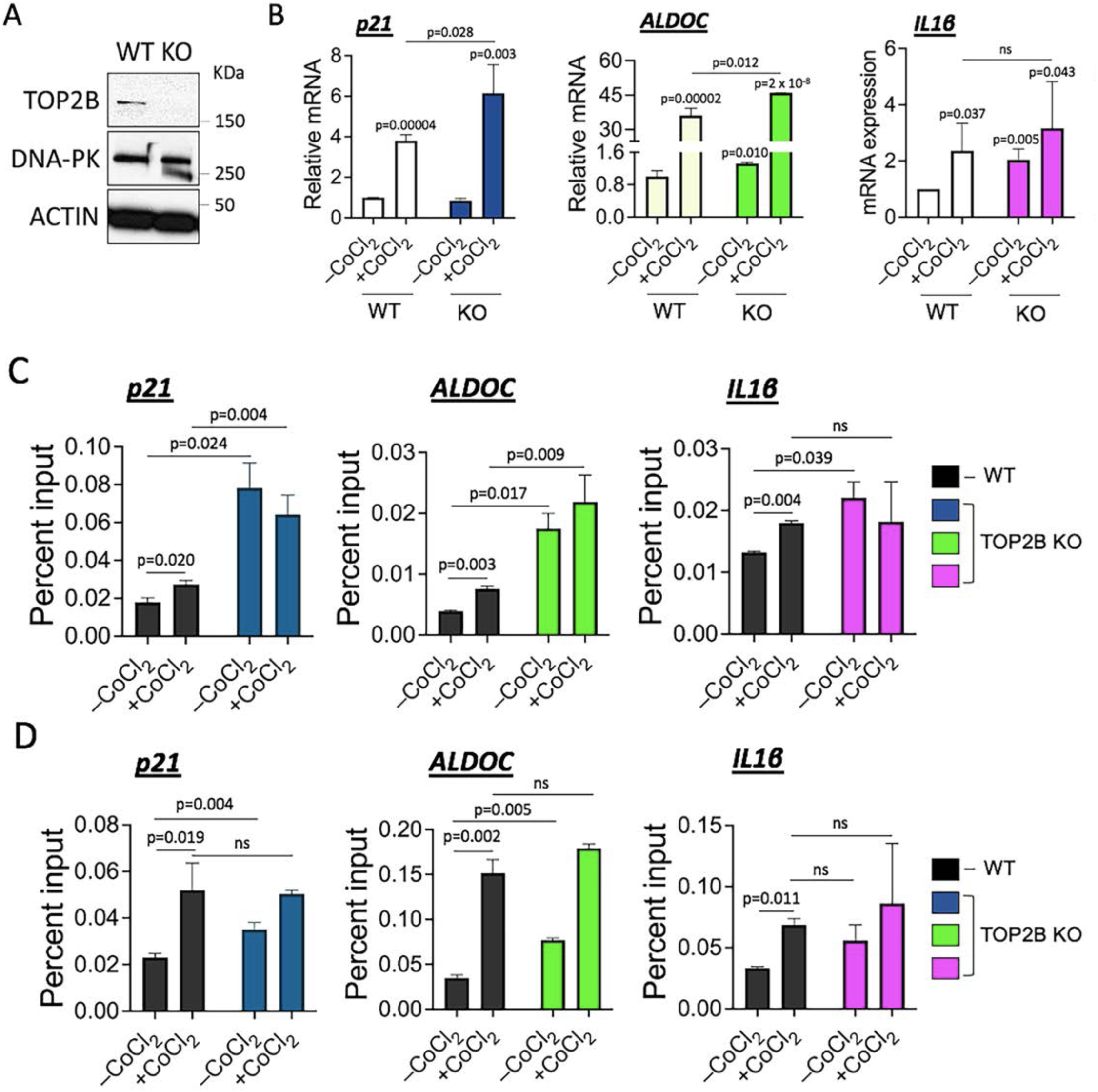
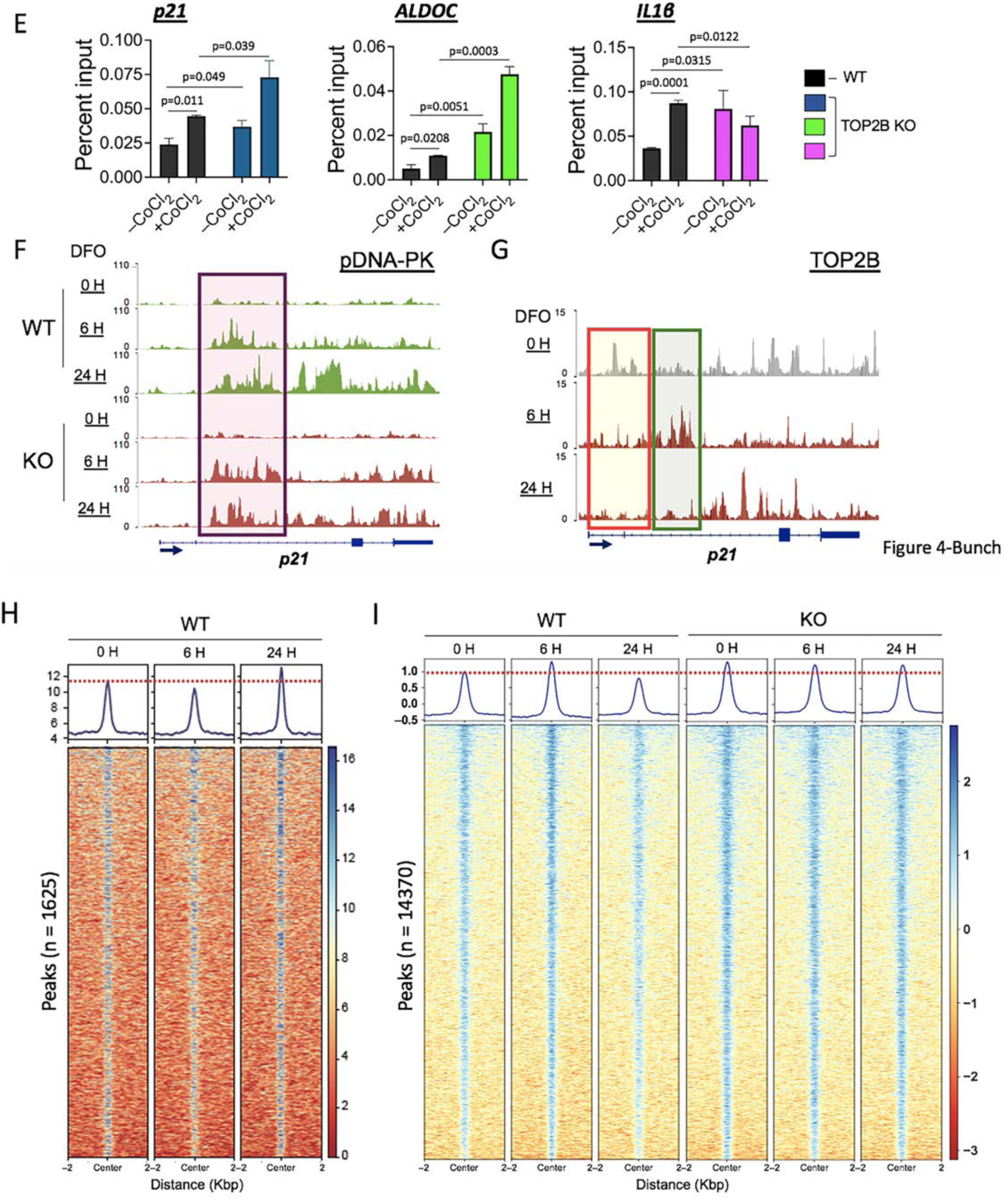
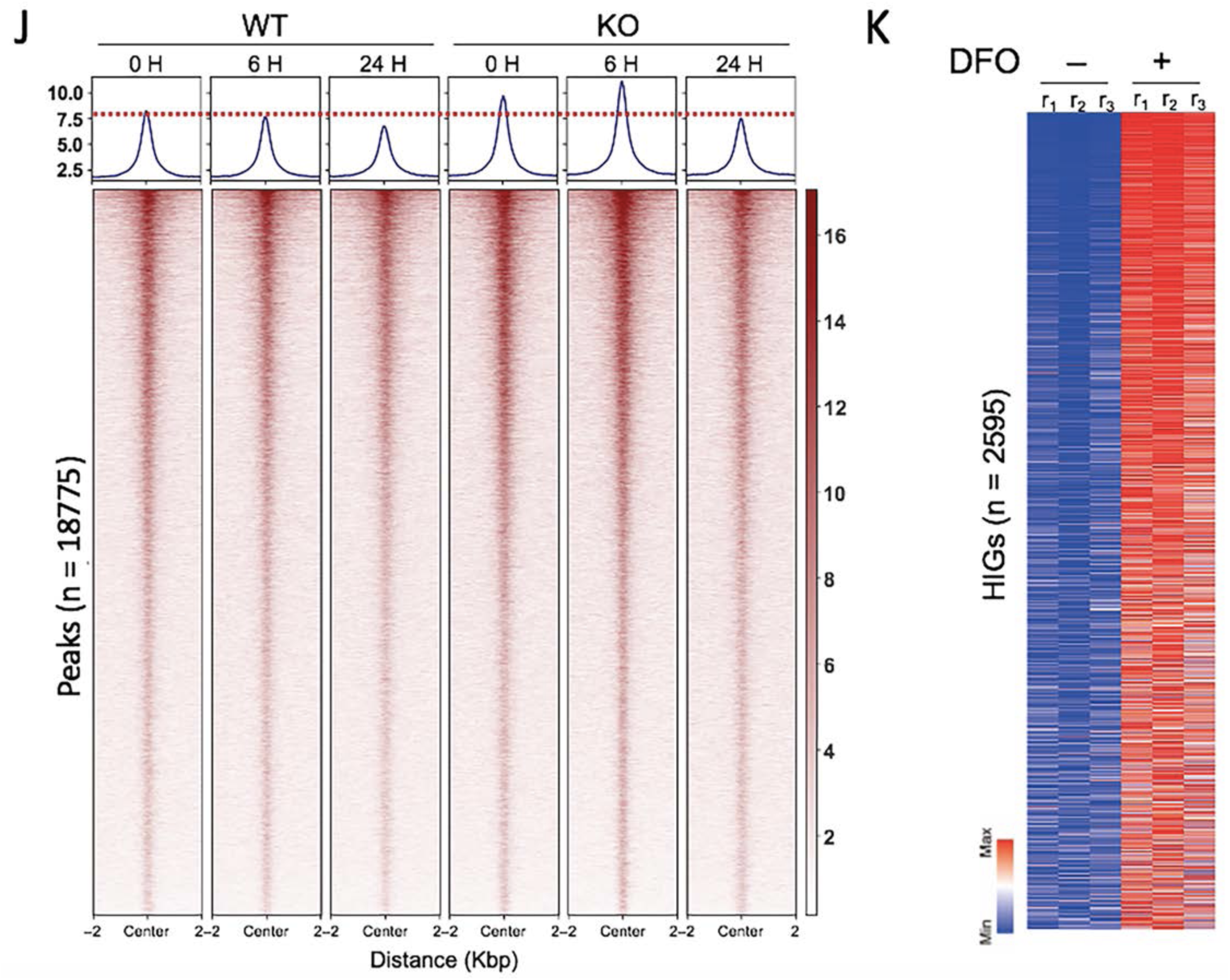
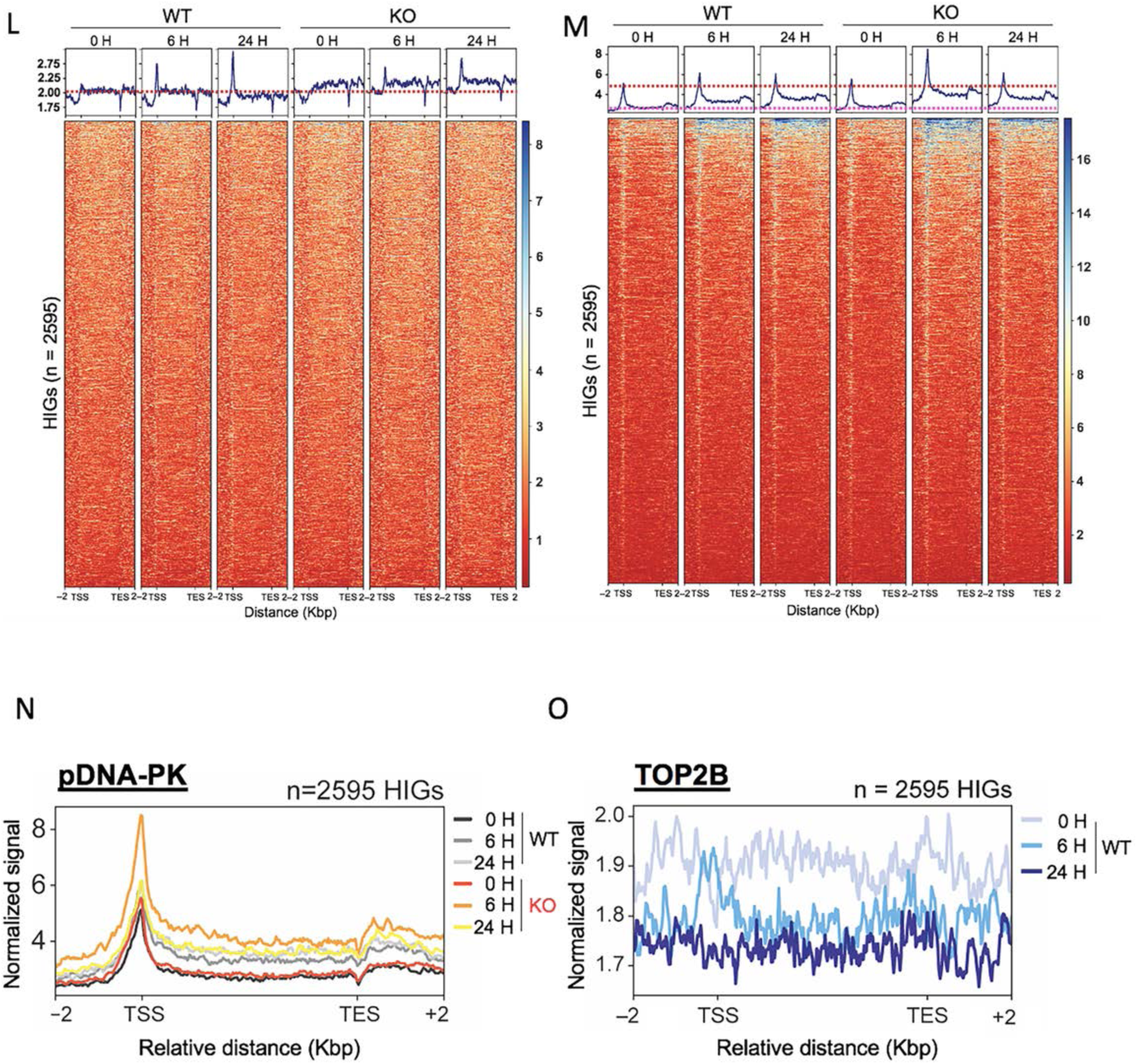
TOP2B regulates HIF1α and DNA-PK recruitment and activity at HIGs. (**A**) Immunoblots showing the protein levels of TOP2B and DNA-PK in WT and TOP2B KO SH-SY5Y cells. ACTIN, a reference. (**B**) qRT-PCR results presenting the effects of TOP2B KO on representative HIG transcription (n = 3 biological samples). Data are presented as mean values and SD. *P*-values for the bar graphs were calculated with the unpaired, one-sided Student’s t-test. (**C**) ChIP-qPCR results showing DNA-PK occupancies on representative HIGs with or without TOP2B (n = 3 independent experiments). Data are presented as mean values and SD. *P*-values for the bar graphs were calculated with the unpaired, two-sided Student’s t-test. (**D**) ChIP-qPCR results showing pDNA-PK occupancies on representative HIGs with or without TOP2B (n = 3 independent experiments). Data are presented as mean values and SD. *P*-values for the bar graphs were calculated with the unpaired, two-sided Student’s t-test. (**E**) ChIP-qPCR results showing HIF1α occupancies on representative HIGs with or without TOP2B (n = 3 independent experiments). Data are presented as mean values and SD. *P*-values for the bar graphs were calculated with the unpaired, one-sided Student’s t-test. (**F**) Chromosome view at *p21* gene showing pDNA-PK occupancies under normoxia and hypoxia for 0 h (normoxia, 0 H), 6 h (6 h hypoxia, 6 H), and 24 h (24 h hypoxia, 24 H) in WT and TOP2B KO (KO) SH-SY5Y cells. Hypoxic stress was induced using 60 μM DFO in **Figs. 4F–O**. A box with purple tint shows notable changes in pDNA-PK recruitment under hypoxic stress and without TOP2B. (**G**) Chromosome view at *p21* gene showing TOP2B occupancies under normoxia and hypoxia for 0 h (normoxia, 0 H), 6 h (6 h hypoxia, 6 H), and 24 h (24 h hypoxia, 24 H) in WT cells. A tinted red box shows TOP2B dissociation from the downsteam of the TSS at p21 gene under hypoxic stress. A tinted green box indicates a possible repositioning of TOP2B in the hypoxic time course. (**H**) Heatmaps from ChIP-seq data showing TOP2B genomic occupancy changes under hypoxic stress in WT SH-SY5Y cells (n = 2 independent experiments). (**I**) Heatmaps from ChIP-seq data showing TOP2B genomic occupancy changes under hypoxic stress in WT and TOP2B KO SH-SY5Y cells (n = 2 independent experiments). (**J**) Heatmaps from ChIP-seq data showing pDNA-PK genomic occupancy changes under hypoxic stress in WT and TOP2B KO SH-SY5Y cells (n = 2 independent experiments). (**K**) RNA-seq data illustrating HIGs (0 h vs 6 h, n = 2595) in SH-SY5Y cells. (**L**) Heatmaps from ChIP-seq results showing HIF1α occupancies in HIGs (n = 2595) under normoxia and hypoxia in WT and TOP2B KO cells (n = 2 independent experiments). (**M**) Heatmaps from ChIP-seq results showing pDNA-PK occupancies in HIGs (n = 2595) under normoxia and hypoxia in WT and TOP2B KO cells (n = 2 independent experiments). (**N**)Metagene plot from ChIP-seq data showing pDNA-PK association on HIGs under normoxia and hypoxia in WT and TOP2B KO cells. (**O**) Metagene plot from ChIP-seq data showing TOP2B occupancies on HIGs under normoxia and hypoxia in WT cells.

ChIP-qPCR analyses under the same conditions demonstrated modest recruitment of DNA-PK to representative HIGs in WT cells under hypoxia (**Fig. 4C**). Interestingly, DNA-PK occupancy was significantly elevated in TOP2B KO cells under normoxic conditions and remained high following hypoxic exposure (**Fig. 4C**). We next assessed phosphorylation of DNA-PK at T2609 (pDNA-PK), a marker of activated DNA-PK which is mediated by autophosphorylation and/or ATM-dependent phosphorylation^23^. In WT cells, pDNA-PK levels increased under hypoxic stress (**Fig. 4D**). Importantly, pDNA-PK occupancy significantly elevated in TOP2B KO cells even under normoxic conditions, consistent with the increased DNA-PK recruitment (**Figs. 4C,D**). For DNA-PK–HIF1α interaction (**Fig. 2A**), we further evaluated HIF1α binding to HIG promoters. As expected, HIF1α was recruited to the genes under hypoxic stress in WT cells (**Fig. 4E**). Notably, HIF1α occupancy was significantly elevated in TOP2B KO cells under normoxic conditions and remained elevated under hypoxia, paralleling the patterns observed for DNA-PK and pDNA-PK, (**Figs. 4C–E**).

To gain insight into genomic TOP2B–DNA-PK-HIF1α dynamics, we performed ChIP-seq analyses in WT and TOP2B KO SH-SY5Y cells following treatment with DFO (or H_2_O for CTRL) to a final concentration of 60 μM for 0, 6, and 24 h (0 H, 6 H, 24 H in figures, respectively). In WT cells, pDNA-PK occupancy increased modestly in the TSS and more prominently in the GB of *p21* under hypoxia (**Fig. 4F**). TOP2B KO increased pDNA-PK levels under normoxic and hypoxic conditions (**Fig. 4F**).

TOP2B occupancy at the *p21* loci was substantially reduced in the TSS under hypoxic stress (**Fig. 4G**). Transient TOP2B enrichment downstream of the TSS at 6 h hypoxia, followed by loss at 24 h, suggested a time-dependent redistribution of TOP2B during hypoxic exposure (**Fig. 4G**). Genome-wide ChIP-seq analysis revealed an initial decrease in TOP2B occupancy at 6 h hypoxia, followed by recovery and further enrichment at 24 h (**Fig. 4H**). These differential patterns were clarified by parallel analysis of HIF1α, which showed reciprocal binding dynamics, with maximal occupancy at 6 h and reduced binding at 24 h (**Fig. 4I**). In TOP2B KO cells, HIF1α occupancy was significantly increased even under normoxia, and remained elevated throughout hypoxic treatment (**Fig. 4I**). Unexpectedly, global pDNA-PK levels were gradually declined in WT cells under hypoxic stress, suggesting attenuation of cellular DNA-PK activity under hypoxia, whereas they were markedly upregulated in TOP2B KO cells under normoxia and hypoxia at 6 h hypoxia (**Fig. 4J**).

We performed RNA-seq analysis in SH-SY5Y cells, comparing the mRNA levels under normoxia and hypoxia (H_2_O control or 60 μM DFO for 6 h). From this approach, 2595 upregulated genes were categorized as HIGs (**Fig. 4K**; **Supplementary Data 5**). HIF1α ChIP-seq analysis revealed increased binding in the TSSs of HIGs under 6 h and 24 h hypoxia in WT cells, consistent with canonical hypoxic transcriptional activation (**Figs. 4E,L**). In TOP2B KO cells, HIF1α occupancy was further increased on these genes, however displayed aberrantly broad distribution across GBs rather than typical, focused enrichment in the TSSs (**Fig. 4L** and **Supplementary Figure 4A**), indicating dysregulated transcriptional control in the absence of TOP2B. Analysis of pDNA-PK ChIP-seq data demonstrated increased occupancy in both TSSs and GBs of HIGs under hypoxia in WT cells (**Figs. 4M**,**N**). Importantly, TOP2B KO markedly amplified pDNA-PK recruitment under both normoxia and hypoxia (**Figs. 4M**,**N**), with particularly strong enrichment at 6 h timepoint (**Figs. 4M**,**N** and **Supplementary Figure 4B**), suggesting a critical regulatory window during hypoxic adaptation involving TOP2B and DNA-PK interaction. Consistent with this, TOP2B occupancy across HIGs was progressively reduced during hypoxia, with a substantial reduction and redistribution at 6 h time point (**Fig. 4O** and **Supplementary Figure 4C**). Together, the data presented in **Fig. 4** support a model in which TOP2B restricts the recruitment and activation of DNA-PK and HIF1α at HIGs and DNA-PK phosphorylation at T2609 and suppresses transcription. By limiting DNA-PK phosphorylation at T2609 and constraining HIF1α binding, TOP2B suppresses inappropriate or excessive HIG transcription. Loss of TOP2B disrupts this regulatory network, leading to hyperactivation of DNA-PK, dysregulated HIF1α occupancy, and enhanced HIG expression.

### TOP2B phosphorylation by DNA-PK is repressive for HIG transcription

Given the physical and functional interactions between TOP2B and DNA-PK (**Figs. 2A,D,F,G,3G,4C,D,F,M–O**), we investigated whether TOP2B is a direct substrate of DNA-PK. Mass spectrometry analysis suggested multiple candidate phosphorylation sites within the CTD of TOP2B (1370–1599 a.a.)(**Supplementary Data 6**). This region is of particular interest because it contains sequences that are highly divergent from TOP2A (**Supplementary Figure 5A**). As we focused on TOP2B in HIG transcription in this study, the phosphorylatable residues within this region were screened via primary sequence comparisons. Five phosphorylatable residues in the TOP2B-unique sequences (underlined, numbering using TOP2B longest isoform 1, **Fig. 5A**) were individually mutated into alanine (A), and the WT and resultant TOP2B mutants were expressed in TOP2B KO SH-SY5Y cells. Although transfection efficiency was low in this cell line, modest but reproducible expression of all constructs were confirmed by immunoblotting (**Fig. 5B**).

**Fig. 5.**
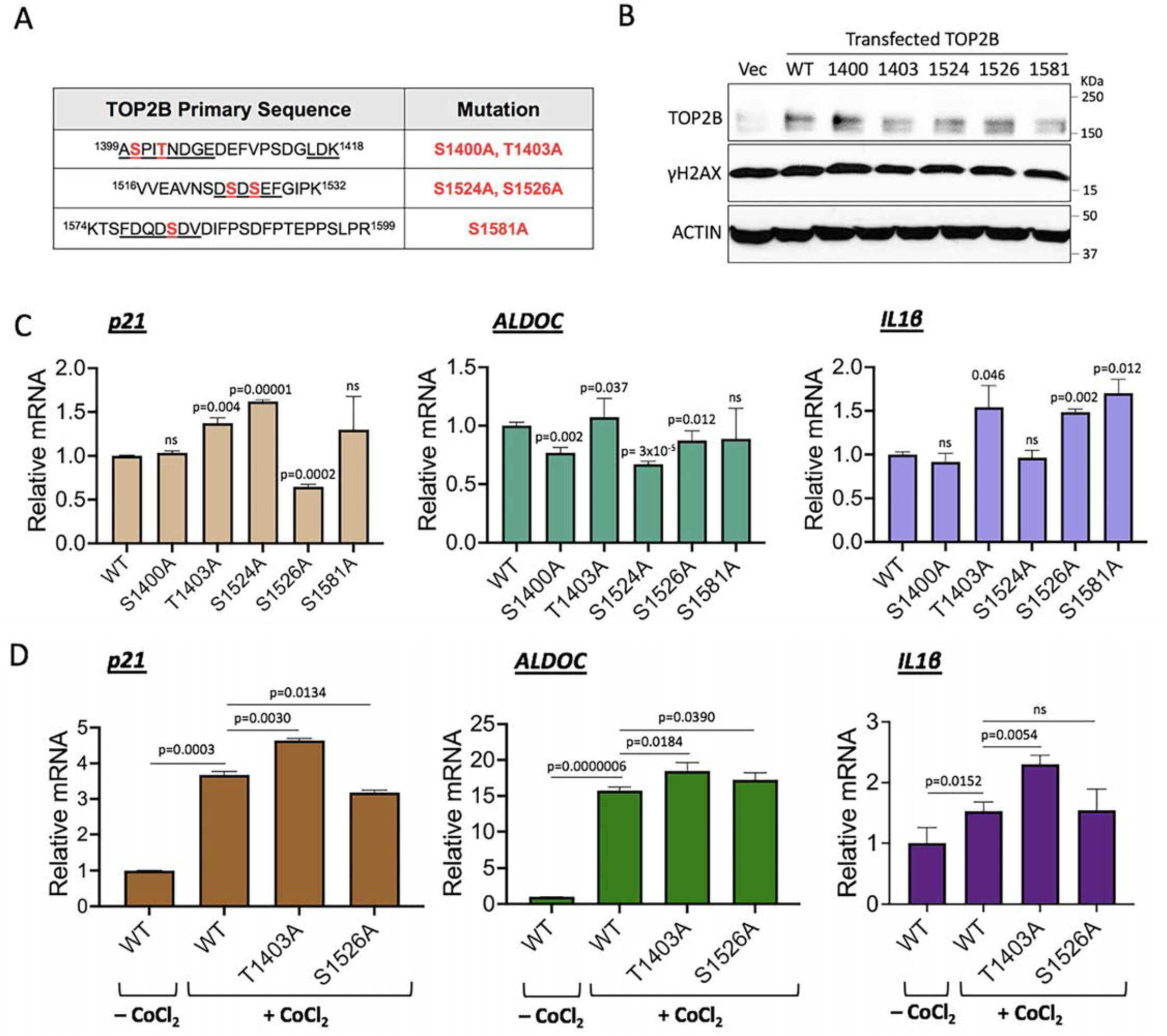
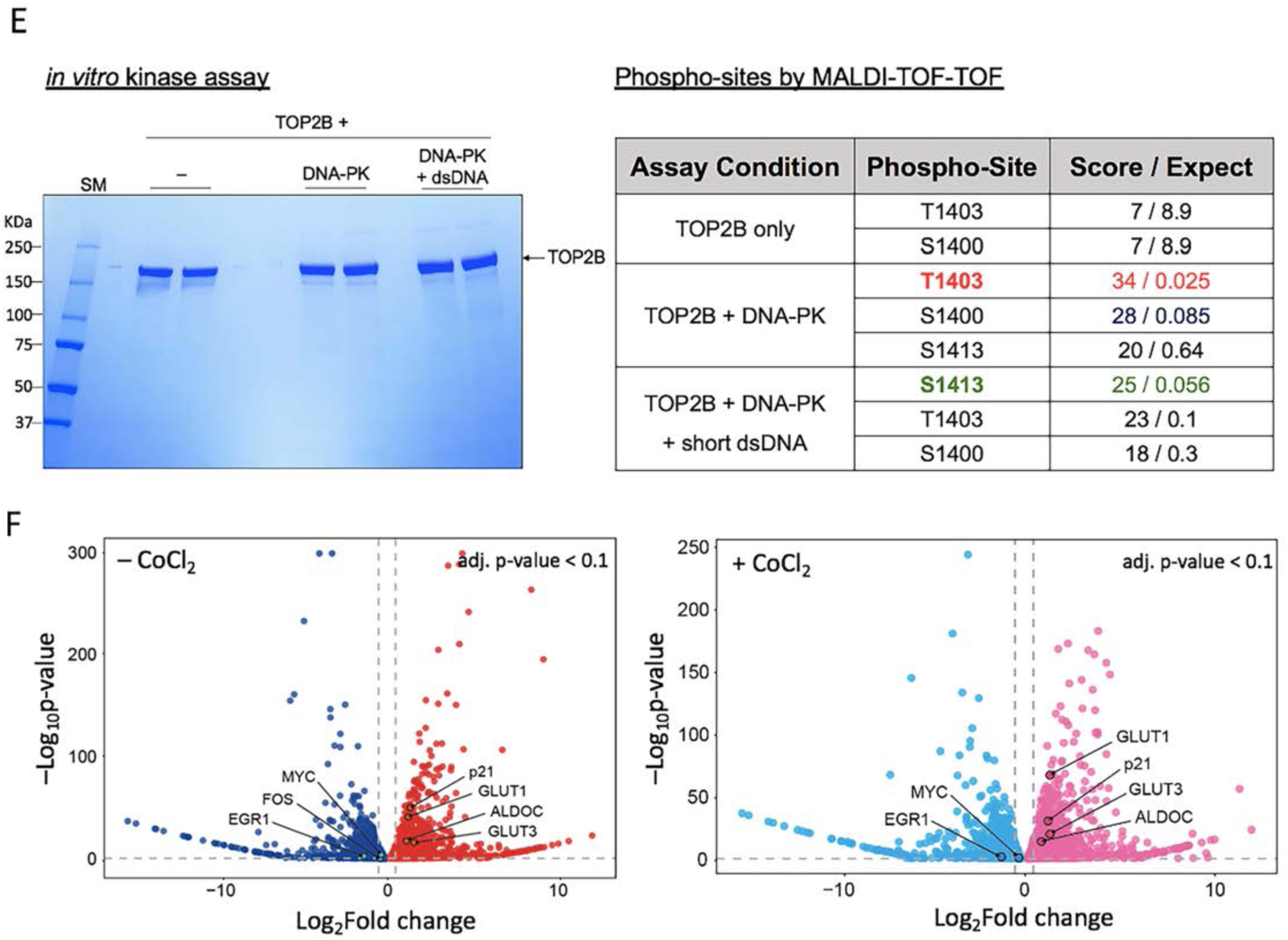
TOP2B phosphorylation by DNA-PK is repressive for HIG transcription. (**A**) Primary sequences of representable TOP2B fragments targeted for mutational analyses in this study. The sequences unique to TOP2B are underlined. The phosphorylatable residues within the unique sequences and mutations generated in this study are in red. (**B**) Immunoblots of TOP2B and γH2AX. Plasmids expressing WT and mutant TOP2B proteins were transfected into TOP2B KO SH-SY5Y cells. Vec, an empty plasmid as a control. ACTIN, a reference. (**C**) qRT-PCR results showing the effects of mutant TOP2B proteins on representative HIG transcription (n = 3). Data are presented as mean values and SD. *P*-values for the bar graphs were calculated with the unpaired, one-sided Student’s t-test. (**D**) qRT-PCR results showing the effects of T1403A and S1526A TOP2B mutations on representative HIG transcription under hypoxic conditions (n = 3). Data are presented as mean values and SD. *P*-values for the bar graphs were calculated with the unpaired, one-sided Student’s t-test. (**E**) Left, a representable coomassie-stained SDS-PAGE gel showing TOP2B protein phosphorylated by DNA-PK in the *in vitro* kinase assay. The gel slices containing TOP2B proteins (180 KDa; marked with an arrow) were subjected to in-gel digestion, followed by mass spectrometry analysis. Right, a table summarizing mass spectrometry data that indicated TOP2B residues, including T1403, phosphorylated by DNA-PK. (**F**) RNA-seq data showing that the expression of a number of genes including representative HIGs, increases, whereas the expression of representative IEGs decreases in DNA-PK KO cells expressing catalytically null DNA-PK mutant (KD), compared to WT cells. Left, normoxia; right, hypoxia.

WT and the five mutant TOP2B-expressing cells were analyzed for the mRNA levels using qRT-PCR. Among these mutants, T1403A consistently increased transcription of the representative HIGs, *p21*, *ALDOC*, and *IL1β* under normoxic conditions. In contrast, the S1526A mutation produced gene-specific effects, decreasing expression of *p21* and *ALDOC* while increasing *IL1β* expression (**Fig. 5C**). These TOP2B mutants were compared with the WT under hypoxic conditions. Cells were supplemented with CoCl_2_ at 200 µM for 12 h at 48 h post-transfection, and the mRNA expression of representative HIGs was quantified using qRT-PCR. Consistent with the data obtained under normoxic conditions, T1403A increased transcription of all three tested HIGs under hypoxia, suggesting that TOP2B phosphorylation at T1403 promotes the transcriptionally repressive function of TOP2B for HIGs (**Figs. 5C,D**). The S1526A mutant again exhibited variable expression patterns for these genes (**Figs. 5C,D**).

To validate TOP2B T1403 phosphorylation by DNA-PK, we performed *in vitro* kinase assays using human TOP2B purified from HEK293 cells and human DNA-PK purified from *C. elegans*, followed by MALDI-TOF-TOF mass spectrometry (**Fig. 5E**). TOP2B protein was deubiquitinated and dephosphorylated during the purification to minimize background modifications^35,36^. The data indicated T1403 as a highest-confidence phosphorylation site targeted by DNA-PK (**Fig. 5F**; **Supplementary Data 7,8**). Interestingly, stimulation of DNA-PK with short dsDNA fragments, mimicking DSBs, shifted phosphorylation preference toward S1413 (**Fig. 5F**; **Supplementary Data 7–9**), suggesting the DNA damage context influences TOP2B phosphorylation by DNA-PK. Consistent with these biochemical findings, RNA-seq analysis comparing WT and KD cells demonstrated that the kinase function of DNA-PK is suppressive to a large-number of genes, including representative HIGs, under both normoxic and hypoxic conditions (**Fig. 5F**; **Supplementary Data 10,11**). In contrast, the transcription of IEGs, including *MYC*, *EGR1*, and *FOS*, is dependent on DNA-PK-mediated phosphorylation (**Fig. 5F**; **Supplementary Data 10,11**), consistent with our previous reports^23,25^. These results highlight gene class-specific regulatory functions of DNA-PK catalytic activity.

To further define the role of TOP2B and TOP2B-DNA-PK interaction in HIG transcription, we performed RNA-seq analysis in WT and TOP2B KO SH-SY5Y cells following 6 h of DFO-induced hypoxic stress. We identified 2595 upregulated and 1836 downregulated genes (**Fig. 6A**; **Supplementary Data 5**). Comparison with HIGs previously identified in HCT116 cells revealed 96 common genes (**Fig. 6B**; **Supplementary Data 12**), which were enriched for carbohydrate metabolisms and HIF1 signaling pathways, which are signature hypoxic responses (**Fig. 6B**). Cell line-specific HIGs were associated with distinct biological pathways, including ferroptosis and mineral absorption for HCT116 cells and p53 and MAPK signaling for SH-SY5Y cells (**Supplementary Figure 6A,B**). These divergences in both the number of HIGs (192 vs 2595 HIGs) and HIG-participating biological pathways likely reflect differences in hypoxia-inducing agents (CoCl_2_ vs DFO) and/or cellular context. TOP2B KO increased and decreased 2278 and 2452 genes, respectively (**Fig. 6C**; **Supplementary Data 13**). It is noteworthy that the genes regulating morphogenesis were most significantly affected by TOP2B KO (**Supplementary Figure 6C,D**) as shown in the TOP2B-bound, DNA-PK KO-increased genes in HCT116 (**Supplementary Figure 3B**). Among the differentially expressed genes, 722 were induced by hypoxia (**Supplementary Figure 6E**; **Supplementary Data 13**).

**Fig. 6.**
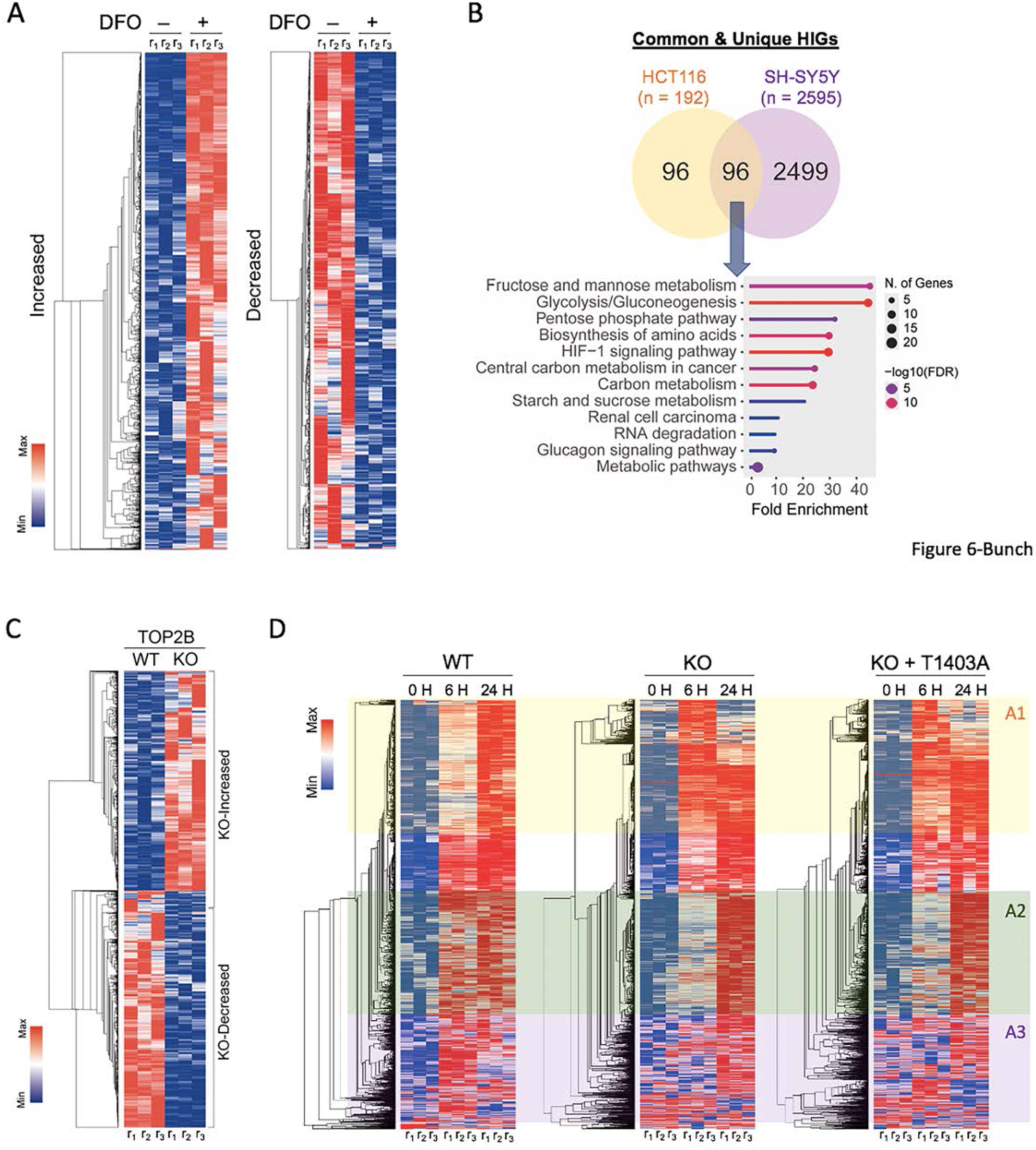
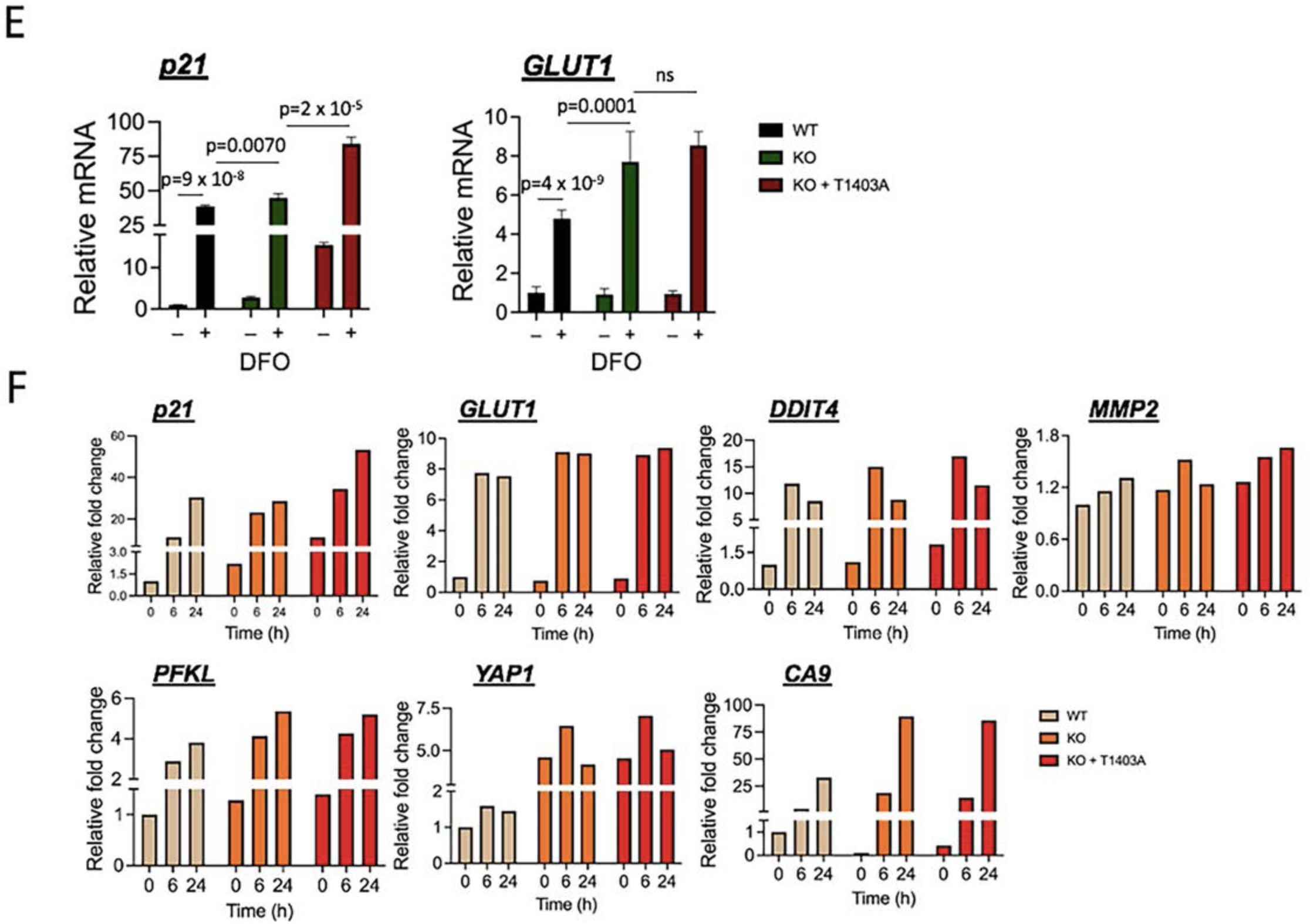
TOP2B controls transcription of a large number of genes including HIGs. (**A**) Heatmaps illustrating HIG expression analyzed by RNA-seq data. Genes, whose expression is significantly increased (left, n = 2595) or decreased (right, n = 1836) under hypoxia induced by 60 μM DFO for 6 h, are shown. |Log_2_FC| > 0.5, p < 0.05. (**B**) Top, Venn diagram comparing HiGs in HCT116 and SH-SY5Y. Bottom, KEGG pathway analysis data with the common 96 HIGs between HCT116 and SH-SY5Y cells. (**C**) Heatmap presenting differentially expressed genes in the absence of TOP2B. Upregulated (n = 2278) and downregulated (n = 2452) genes. |Log_2_FC| > 0.5, p < 0.05. (**D**) Heatmaps showing the expression changes of 2595 HIGs under normoxia and hypoxia in WT, TOP2B KO, and TOP2B KO expressing T1403A TOP2B (KO + T1403A, hereafter). Area 1 (A1, yellow) indicates the genes whose expression is increased under 6 h hypoxic stress (6 H) in KO and KO + T1403 cells, compared to WT cells. A2 (green) indicates the genes whose expression is increased under 24 h hypoxic stress (24 H) in KO and KO + T1403A cells, compared to WT cells. The expression of those genes marked with A3 (purple) upregulated under normoxia in KO and KO + T1403A cells, compared to WT cells. (**E**) qRT-PCR results showing the effects of TOP2B KO and T1403 mutation on representative HIGs, *p21* and *GLUT1* transcription (n = 3). Data are presented as mean values and SD. *P*-values for the bar graphs were calculated with the unpaired, one-sided Student’s t-test. (**F**) mRNA expression of representative HIGs from RNA-seq data. Relative fold changes under nomoxia (0 h) and hypoxia (6 h and 24 h) in WT (tan), KO (orange), and KO + T1403A (red) cells. p < 0.05.

We next compared mRNA levels of the 2595 HIGs in WT, TOP2B KO, and TOP2B KO expressing the phospho-null T1403A TOP2B mutant. RNA-seq analysis showed that the majority of HIGs were increased in TOP2B KO and T1403A-expressing cells, compared to WT cells. These genes were shown in three categories, depending on the time-point of increased gene expression: 6 h (A1, marked yellow), 24 h (A2, marked green), or 0 h (normoxia, A3, marked purple)(**Fig. 6D**). These genome-wide findings were validated by qRT-PCR and RNA-seq analyses of representative HIG genes, including *p21*, *GLUT1*, *MMP2*, *DDIT4*, and *YAP1*. Loss of TOP2B significantly increased expression of these genes, and reconstitution with the T1403A mutant further potentiated the increases (**Figs. 6E**,**F**). Overall, the data presented in **Figs. 5** and **6** support a model in which DNA-PK-mediated phosphorylation of TOP2B at T1403 is critical for the suppressive function of TOP2B in HIG transcription.

### TOP2B restricts “DNA openness” to prevent HIG activation under normoxia

To elucidate the mechanism of DNA-PK-mediated TOP2B regulation, we employed trimethylpsoralen–sequencing (TMP–seq). TMP is a small organic chemical that preferentially intercalates into negatively supercoiled or underwound DNA and TMP-DNA complexes can be covalently cross-linked by UV light^69,70^. The genomic regions incorporated by TMP were enriched by removal of TMP-free DNA regions through chromatin fragmentation and repeated DNA denaturation and digestion using DNases (**Fig. 7A**). We first compared DNA topological changes in WT and TOP2B KO SH-SY5Y cells. Our metagene analyses showed that genomic regions (n = 60649 genes; –1 Kb from TSS to +1 Kb from transcription end site [TES]) were markedly underwound in TOP2B KO cells, compared with WT cells (left panels, **Fig. 7B**). Consistently, open DNA regions surrounding TSSs (n = 60649 genes; –0.5 Kb to +1 Kb from TSS) were more prominent in TOP2B KO cells than WT cells (heatmaps on the right panels, **Fig. 7B**). We next examined DNA topology at the genes that were identified as hypoxia-induced (n = 192) or -repressed (n = 66) by RNA-seq (**Figs. 3B,D,F**) under normoxia and hypoxia (**Fig. 7C** and **Supplementary Figures 7A,B**). Strikingly, DNA around approximately +200–500 regions became markedly negatively-supercoiled or underwound under hypoxia-inducible gene activation (marked with a red asterisk; **Fig. 7C**). In contrast, this spike of open DNA topology formation under hypoxia was not observed in TOP2B KO cells, although DNA were overall more open near the TSSs under normoxia (marked with yellow tint; **Fig. 7D**). To understand the relation between DNA underwinding/negative supercoiling and chromatin accessibility, we examined the 192 HIGs through micrococcal nuclease (MNase)-seq under identical conditions to those used for TMP-seq analysis. MNase-seq revealed that hypoxia increased overall accessibility at HIGs and identified specific regulatory regions that were differentially accessible under normoxic and hypoxic conditions (**Fig. 7E**; marked with blush tint). Importantly, these regions were closely overlapped with the topological hot spots identified by TMP-seq (**Figs. 7C,E**). For example, the +200–500 DNA region exhibited both markedly increased underwinding/negatively supercoiling and dramatically enhanced accessibility (**Figs. 7C,E**) under hypoxic stress. In contrast, the –500 promoter region was already open and accessible under normoxia and became further underwound and accessible under hypoxia (**Figs. 7C,E**). In TOP2B KO cells, chromatin accessibility was elevated even under normoxia (**Fig. 7F**), consistent with increased basal DNA underwinding shown in **Fig. 7D**. However, the two regulation hot spots identified in WT cells were significantly deregulated in the absence of TOP2B (**Figs. 7C–F**). Interestingly, hypoxic stress reduced DNA accessibility in TOP2B KO cells (**Fig. 7F**), suggesting impaired adaptive chromatin remodeling.

**Fig. 7.**
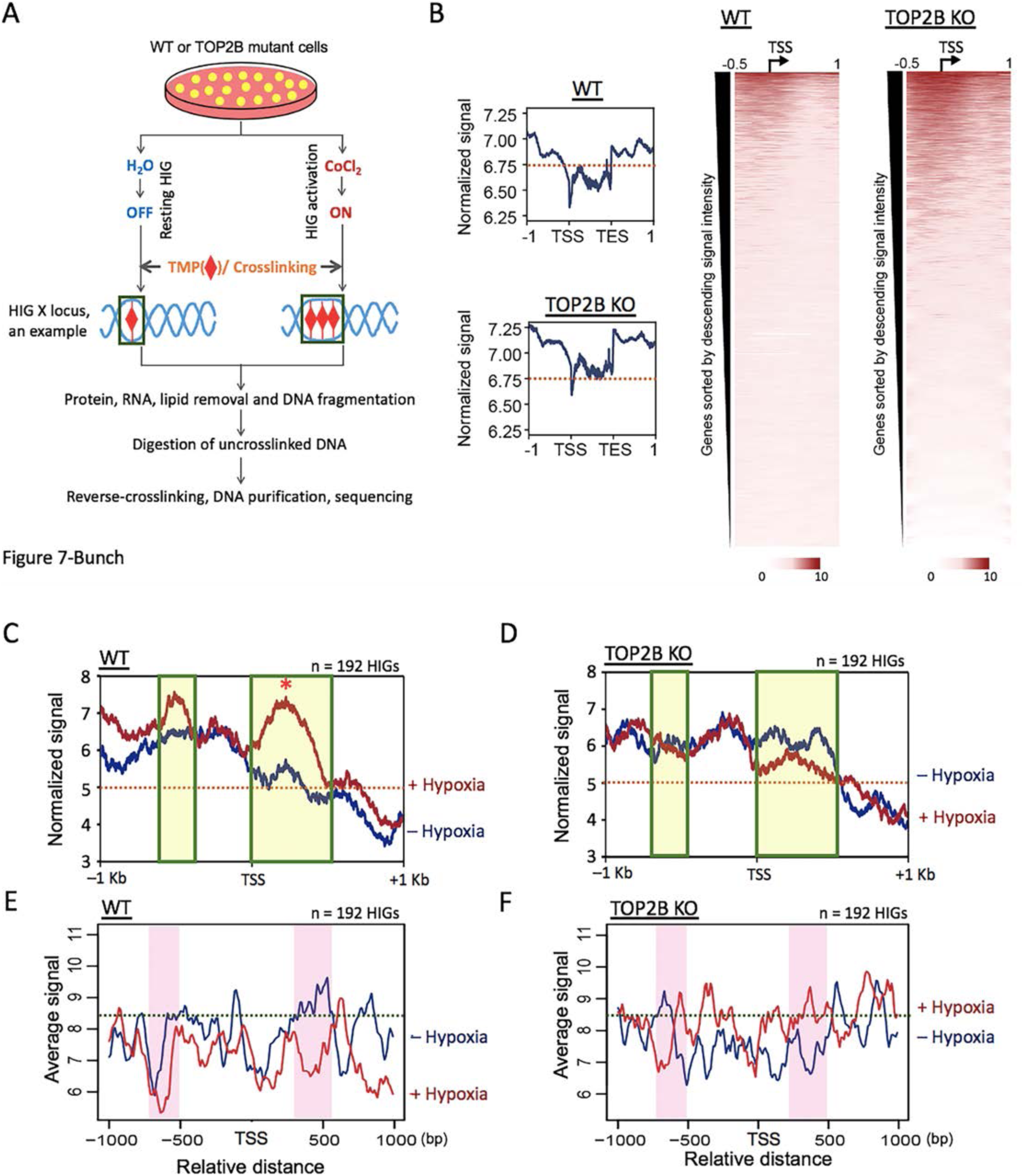
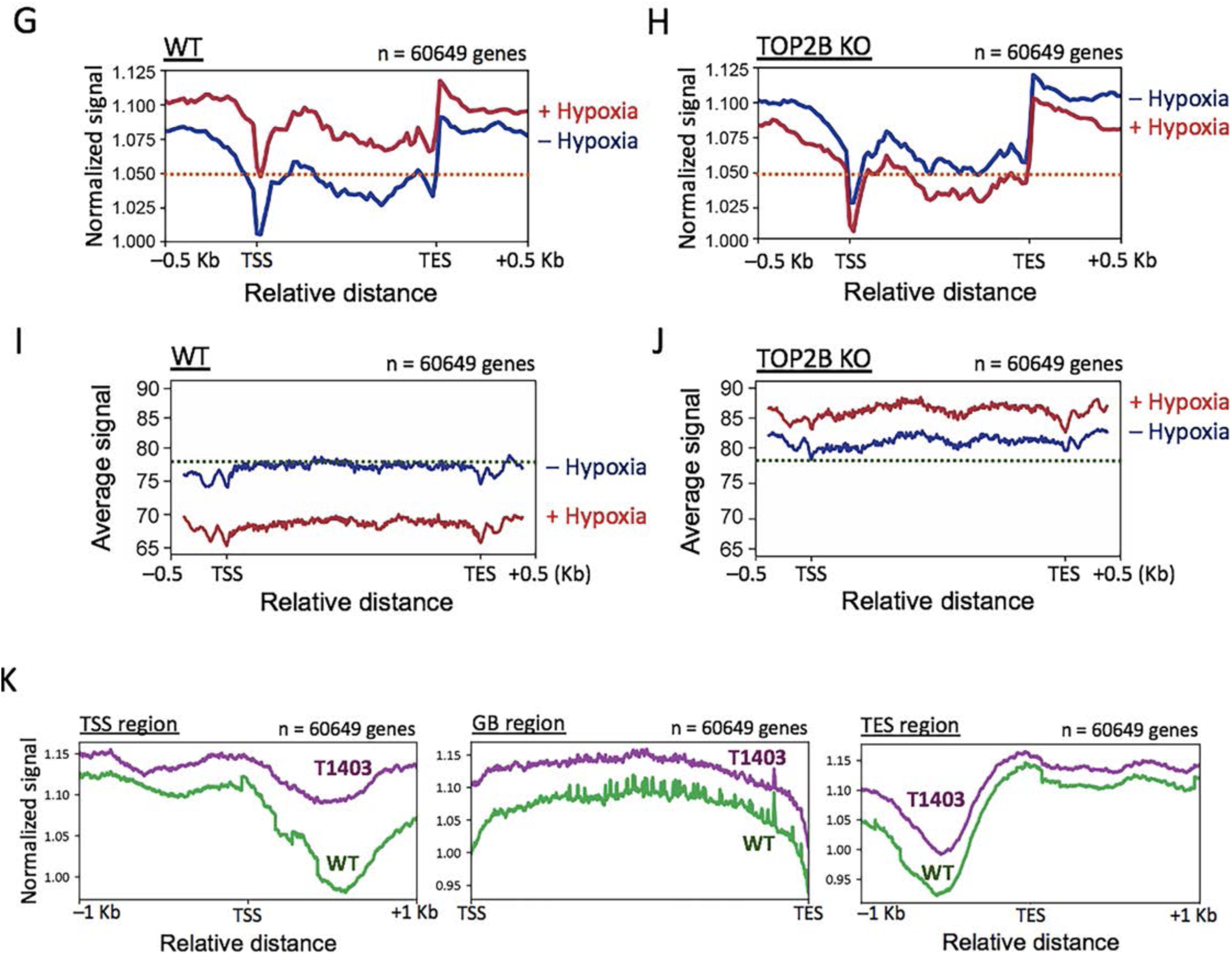
TOP2B restricts “DNA openness” to prevent HIG activation under normoxia. (**A**) Schematic representation of TMP-seq sample preparation. TMP, red diamond; cells, yellow circles; DNA, double sky-blue lines. (**B**) Metagene analyses (left, line graphs) and heat maps (right), comparing underwound genomic regions in WT and TOP2B KO cells (n = 2 independent experiments). X axis shows distance from TSS or TES in Kilobases. (**C**) Metagene analyses from TMP-seq showing DNA topology changes in HIGs (n = 192) under normoxia (–hypoxia) and hypoxia (+hypoxia) in WT cells. The genomic region undergoing dramatic DNA unwinding under hypoxia (approximately +200–500 from TSS) is marked with a red asterisk in a green box. (**D**) Metagene analyses from TMP-seq showing DNA topology changes in HIGs (n = 192) under normoxia (– Hypoxia) and hypoxia (+ Hypoxia) in TOP2B KO cells. DNA is more underwound under normoxia and the dramatic DNA unwinding under hypoxia in WT cells, marked in a green box as regulation hot spots, is deregulated in the absence of TOP2B. (**E**) Metagene analyses from MNase-seq showing DNA accessibility changes in HIGs (n = 192) under normoxia (– Hypoxia) and hypoxia (+ Hypoxia) in WT cells (n = 2 independent experiments). The regions correlative to hot spots for DNA topological regulations are marked in blush. These regions become accessible under hypoxia. (**F**) Metagene analyses from MNase-seq showing increased overall DNA accessibility under normoxia and significant DNA accessibility deregulation under hypoxia in HIGs (n = 192) in TOP2B KO cells. (**G**) Metagene analyses from TMP-seq of all genes (n = 60649) in WT. Hypoxia increases DNA underwinding/negative supercoiling. (**H**) Metagene analyses from TMP-seq of all genes (n = 60649) in TOP2B KO cells. Without TOP2B, genomic regions are more underwound under normoxia and proper hypoxia-mediated gene induction is deregulated. (**I**) Metagene analyses from MNase-seq of all genes (n = 60649) in WT. Hypoxia increases DNA accessibility. (**J**) Metagene analyses from MNase-seq of all genes (n = 60649) in TOP2B KO cells. Without TOP2B, genomic regions become less accessible under normoxia, hypoxia further decreases DNA accessibility genome-wide. (**K**) Metagene analyses from TMP-seq of all genes (n = 60649) in TOP2B KO cells transfected with WT TOP2B (sky-blue line)- or T1403A TOP2B (red line)-expressing plasmids. TOP2B T1403A, a mutation unphosphorylatable by DNA-PK, increases DNA underwinding throughout the promoter, TSS, GB, and TES, in particular at around +500 regions from the TSS, genome-wide.

Genome-wide metagene analyses (n = 60649) showed the architecture of genome, in which GB regions are overall less open than promoters or TES-flanking regions under normoxia (**Fig. 7G**; – hypoxia). Hypoxia induced widespread DNA unwinding across promoters, TSSs, GBs, and TESs in WT cells (**Fig. 7G**; + hypoxia). In contrast, TOP2B KO cells exhibited extensive genomic openness/negative supercoiling under normoxia (**Fig. 7H**; – hypoxia), without additional DNA underwinding but more winding, under hypoxia (**Fig. 7H**; + hypoxia). In WT cells, hypoxia-induced genomic unwinding (**Fig. 7G**) correlated with hypoxia-enhanced genomic accessibility, revealed by MNase-seq analysis (**Fig. 7I**). Notably, while TOP2B KO increased genomic accessibility on HIGs (**Fig. 7F**), it markedly reduced genomic accessibility and further decreased it under hypoxic stress (**Fig. 7J**). These data indicate that TOP2B functions to suppress DNA openness and accessibility at HIGs under normoxia, while enhancing overall genomic accessibility, through modifying DNA winding To understand the role of DNA-PK-mediated TOP2B phosphorylation in genomic DNA topology, the WT TOP2B and the TOP2B T1403A mutant, which showed a consistent increase in the transcription of representative HIGs (**Figs. 4B,5C,D,6C–F**), were expressed in TOP2B KO cells, and the underwound DNA regions under hypoxia were mapped through TMP–seq. It was notable that a valley, presumably more tightly wound DNA regions, located at approximately +500 almost disappeared in T1403A TOP2B cells (**Fig. 7K**, left panel), suggesting that TOP2B phosphorylation at T1403 by DNA-PK is responsible for relaxing the negative supercoiling formed in the downstream of TSSs. Moreover, GB and TES accumulated more open DNA structures in T1403A TOP2B cells, compared with WT cells (middle and right panels, **Fig. 7K**). These open DNA topological changes were consistent with more active transcription of representative HIGs in TOP2B T1403A and DNA-PK KO/KD cells (**Figs. 2D,3A,C–G,5C,D,F,6D–F**), suggesting that DNA-PK-mediated phosphorylation of TOP2B at T1403 enables controlled transcription by suppressing negative supercoiling/underwound DNA formation in HIGs.

## DISCUSSION

In this study, we identified TOP2B as a novel factor to regulate human HIG transcription, and demonstrated that TOP2B dynamically interacts with DNA-PK. These molecules appear to regulate the activities of each other for inducible HIG transcription. TOP2B has been widely characterized as a transcriptional activator of diverse gene classes, including *HSP70*, IEGs, androgen/estrogen receptor genes, and neuronal activity-inducible genes as shown in our previous work and that of others^22,23,25,30,33–36^. Therefore, our finding that TOP2B functions as a transcriptional repressor of HIGs is unexpected and highlights an unappreciated genomic context-dependent role for this enzyme. In addition, our mutational and RNA-seq analyses suggest that DNA-PK plays dual roles on HIG transcription, as an activator for some and a repressor for other HIGs (**Fig. 3** and **Supplementary Figure 3** and **Supplementary Data 1,2**), rather than an activator for HIGs as previously reported^27^.

Our data support the concept that DNA-PK is important for HIF1α recruitment as previously reported^27^ and suggest that DNA-PK KO devastates HIF1α stabilization under hypoxia (**Figs. 2C,F** and **Supplementary Figure 2A,B**). Interestingly, however, DNA-PK deficiency markedly compromises HIF1α recruitment, yet HIG transcription remains robust in DNA-PK KO cells (**Figs. 3A–F** and **Supplementary Figure 3A,B**). This may be explained by a dependence on or a transition to HIF2 in HIF1 instability^71,72^, and that DNA-PK has a suppressive role in the transcription of many HIGs through its interaction with TOP2B.

Our data suggest that TOP2B associates with HIGs and prevents DNA-PK and HIF1α from being recruited under normoxia (**Figs. 4A–O**). DNA-PK recruitment under hypoxia is likely to be important for TOP2B release from the HIGs as DNA-PK KO overturns TOP2B release (**Figs. 2D,G**). Moreover, DNA-PK presence also controls the ability of TOP2B activity to be suppressive because, without DNA-PK, TOP2B stimulates HIG transcription without being released under hypoxia and even under normoxia (**Figs. 2D,E,G,3A–G**). An unphosphorylatable mutant TOP2B, T1403A and genomic topology/accessibility studies suggest that TOP2B removes negative supercoiling formation and reduce DNA accessibility to suppress HIG transcription, and DNA-PK phosphorylation presumably stimulates TOP2B catalysis to relax negative DNA supercoiling (**Figs. 5A–F,6D–F,7B–K**). It is notable that normal hypoxic responses release TOP2B (**Figs. 1,2F,4G,H,O**) and involve a dramatic increase in negative supercoiling or DNA underwinding in the TSSs of HIGs, which was only observed in the WT, not in TOP2B KO cells (**Figs. 7B–F**). These observations in HIG transcription may be in agreement with a previous study showing that transcriptionally-active regions in the genome are overall underwound and tend to be devoid of TOP2^73^.

In HIG regulation, both TOP2A and TOP2B appear to play important roles under normoxic conditions (**Figs. 1B–E** and **Supplementary Figure 1B,C**) to maintain the resting state of HIG expression.

Although both TOP2A and TOP2B dissociate from HIGs under hypoxic stress (**Figs. 1B,C**), the degree and intensity of their association with chromosomes are apparently somewhat distinctive (**Fig. 1D** and **Supplementary Figure 1C**). In addition, previous studies^23,24,33–36,59,74^ suggest mutual and/or distinctive functions of TOP2A and TOP2B in gene regulation. While elucidating the precise contribution of TOP2A to HiG transcription requires further investigations, the present study focuses mainly on understanding the novel function of TOP2B in HIG regulation. In our model (**Fig. 8** and **Supplementary Figure 1C**), we propose that TOP2B relaxes negative DNA supercoiling to prevent it from accumulating and restricts DNA accessibility in the TSSs and also interferes with DNA-PK and HIF1α recruitment to HIG promoters under normoxia. Under hypoxia, DNA-PK recruitment and activation, along with HIF1α, presumably mediates TOP2B release and activates transcription. Our current data suggest that DNA-PK phosphorylates TOP2B at T1403, which stimulates TOP2B catalytic activity to resolve negative supercoiling under normoxia (**Figs. 5A–F,6D–F,7K**). In addition, in the absence of DNA-PK, TOP2B recruitment and activity are augmented under normoxia and hypoxia (**Figs. 2D,E,G,H**). These observations suggest that TOP2B without DNA-PK might be catalytically ineffective to resolve negative supercoiling, despite prolonging the retention time, thus is compromised to suppress gene expression, and/or might have other ways to promote transcription, such as providing protein-protein interactions and structural constituents.

**Fig. 8.**
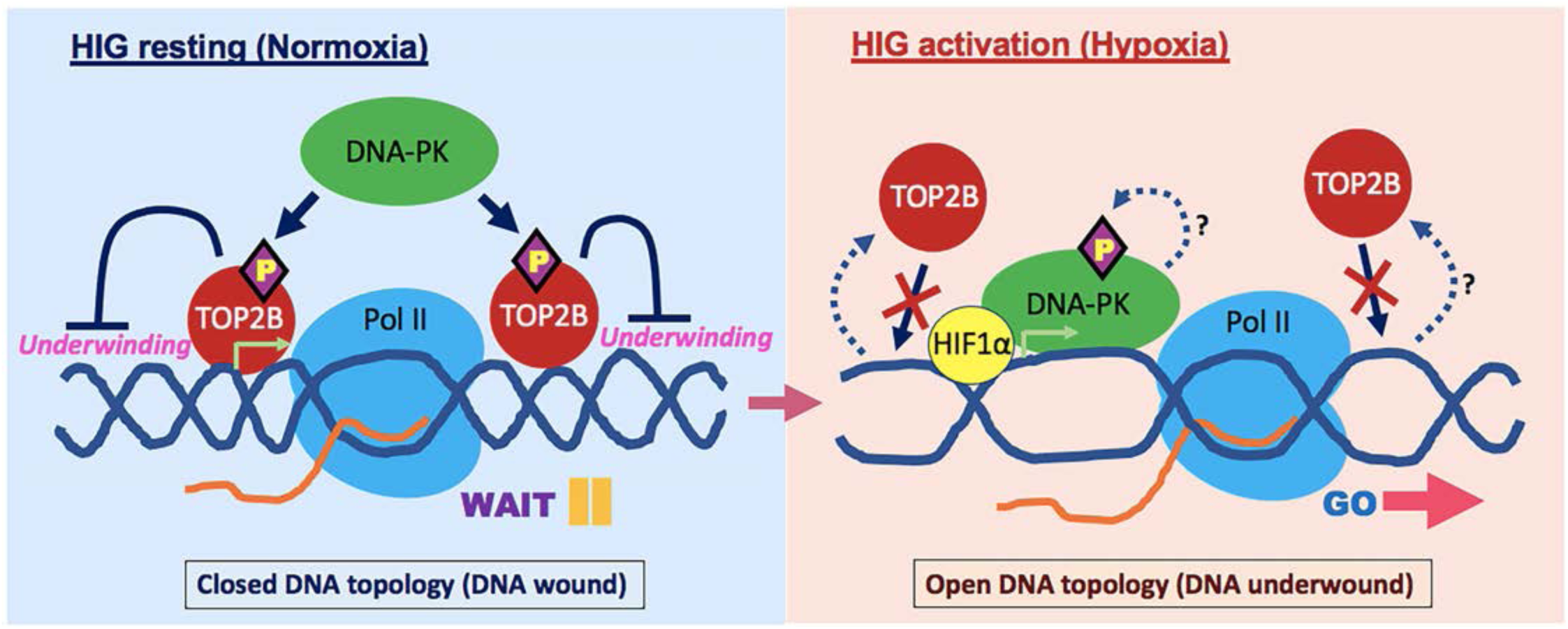
A model of TOP2B regulation in HIG transcription. Under normoxia, TOP2B is associated with HIGs and DNA-PK phosphorylates TOP2B at the CTD including T1403 (phosphorylation marked with a yellow letter “P” in a purple diamond). This phosphorylation is likely to stimulate TOP2B to relax negative DNA supercoiling, preventing DNA underwinding and gene activation. Under hypoxia, TOP2B is released from HIGs by currently unknown mechanisms. Instead, DNA-PK is recruited to HIGs along with HIF1α and is phosphorylated at T2609. Nascent RNA, an orange curved line. TOP2B release and inactivation promote DNA openness/underwinding to facilitate productive Pol II transcription in HIGs. Less defined and characterized processes are shown in dashed lines with question marks. DNA, double blue helical lines; RNA, orange curved lines; TSS, a light green arrow on the DNA.

It is unclear how TOP2B is released from HIGs and what the exact function of DNA-PK, except for augmenting HIF1α recruitment, is under hypoxia, a question which awaits further investigations. It is particularly of interest to ask whether the role of DNA-PK involves DNA repair or some other aspect of HIG activation. DNA-PK appears to switch functions depending on the DNA structures that it associates with. In a recent study, for example, identified that binding to the open DNA ends, DNA-PK suppresses autophosphorylation at T2609 but phosphorylates other protein targets^75^. Another previous study reported unique phenomena, particularly that TOP2B is recruited to the genome loci of DSBs^76^. It is possible that DNA break causes underwound DNA and topological stresses, requiring TOP2B to restore the superhelicity. Also, it is plausible that TOP2B recognizes the broken ends after DNA break as intermediate (transition state) substrates as TOP2B binds to twisted DNA, induces DNA break, and joins the DNA ends, suggesting that TOP2B binds to both intact and broken DNA ends. In the latter case, Ku proteins and DNA-PK may substitute TOP2B on the lesion upon the release of TOP2B in HIG activation. The resolution of these conjectures and the revelation of molecular events and mechanisms including the local DNA structure changes, in which TOP2B and DNA-PK are involved for gene regulation, will be a goal for future studies.

Our previous study showed that TOP2B ubiquitination by the BRCA1-BARD1 complex is important for TOP2B-*EGR1* TSS interaction during transcriptional rest. Upon transcriptional activation, the phosphorylation of BRCA1 at S1524 by ATM/ATR discouraged the BRCA1-BARD1-mediated TOP2B ubiquitination to lower the affinity between TOP2B and *EGR1* DNA^35^. In addition, ERK2 phosphorylates the CTD of TOP2B and regulates its catalytic activity to resolve the positive DNA supercoiling^36^. On the other hand, ERK2 delays the TOP2B catalysis of relaxing the negative DNA supercoiling^36^. In this study, DNA-PK phosphorylates TOP2B to augment the catalysis of negative DNA supercoiling, an event disfavoring productive transcription of HIGs under normoxia. These findings indicate that post-translational modifications of TOP2B by gene-specific transcription factors contribute to the modulation of TOP2B: TOP2B binding to and catalysis of its substrate DNA and TOP2B release from the resulting (product) DNA. Currently, the CTD of TOP2B, which is the hub of numerous modifications including acetylation, phosphorylation, methylation, sumoylation and ubiquitination etc. (https://www.phosphosite.org/proteinAction.action?id=5866&showAllSites=true), has not been structurally resolved, meaning that this domain is highly flexible, unstructured from the core domains^36^. It also suggests that TOP2B CTD composed of approximately 400 amino acids is available to interact with various transcription factors for the stepwise regulation of this enzyme. DNA is the paper for Pol II to copy and the road for Pol II to run and the DNA topology is key regulator for Pol II processivity and productivity. We propose that DNA-PK-mediated TOP2B phosphorylation and regulation in HIG transcription is an important, gene-specific topological element in Pol II transcription.

## Acknowledgments

We are grateful to C. A. Austin at Newcastle University in the United Kingdom for sharing the TOP2B KO SH-SY5Y cells and to D. J. Taatjes at the University of Colorado at Boulder for sharing the HeLa nuclear extracts. We also thank Omega Bioservices and Azenta Life Sciences in the USA and Rokit Genomics in Korea for providing next generation sequencing services. We appreciate Genomine, Inc. in Korea for mass spectrometry analyses, and M. Seu, S. Ju, J. Jeong, and the current members of the Bunch laboratory members at KNU for their technical assistance. H.B. thanks J. N. Park, S. W. Yang, John and D. Y. Bunch and J. Christ for their loving encouragement and support throughout the course of this work. This research was supported by grants from Thomas H. and Dorthy S. Corson Career Development Award in Cardiovascular Disease Research, funded by the Center for Clinical and Translational Science, Mayo Clinic to M.J.S., from the Korea Health Technology R&D Project through Korea Health Industry Development Institute (KHIDI), funded by the Ministry of Health & Welfare of the Republic of Korea (R5-2024-00437643) to J.J., from the National Institutes of Health (R0CA276058 & R01CA29290), the Department of Energy (DE-SC0025578), and the National Aeronautics and Space Administration (80NSSC23K1018) to A.J.D., and from the National Research Foundation (NRF) of the Republic of Korea (2022R1A21003569, RS-2025-02303149, and RS-2025-16067324) to H.B.

## Author Contributions

HB and JL performed ChIP-PCR/qPCR and qRT-PCR. JP, J-YJ, IJ, and KK carried out bioinformatics analyses. JS and M(J)S designed, cloned, and prepared WT and mutant TOP2B plasmids for transfection. HL prepared for and cultured DNA-PK KO and mutant HCT116 cells and provided discussions for proteomics analyses. HB performed *in vitro* kinase assay, ICE assay, immunoprecipitation, and immunoblotting. HB and J-SY carried out mammalian transfection. HB prepared for the samples for mass spectrometry, RNA-seq, ChIP-seq, MNase-seq, and TMP-seq. JJ, YY, SC, KK, and HB performed the genomic data analysis and revised the manuscript. HB created the hypothesis, designed, coordinated the experiments, and wrote the manuscript. AD and HB analyzed and curated the data, revised the manuscript, and obtained the major grants to support this study.

## Competing Interests Statement

The authors declare that they have no competing interests.

## Notes

### Competing Interest Statement

The authors have declared no competing interest.

### Summary of Updates

More experiments have been performed and the data have been newly included in the revision.

